# Diel Investments in Phytoplankton Metabolite Production Influenced by Associated Heterotrophic Bacteria

**DOI:** 10.1101/2020.11.18.388827

**Authors:** Mario Uchimiya, William Schroer, Malin Olofsson, Arthur S. Edison, Mary Ann Moran

## Abstract

Organic carbon transfer between photoautotrophic and heterotrophic microbes in the surface ocean mediated through metabolites dissolved in seawater is a central but poorly understood process in the global carbon cycle. In a synthetic microbial community in which diatom extracellular release of organic molecules sustained growth of a co-cultured bacterium, metabolite transfer was assessed over two diel cycles based on per cell quantification of phytoplankton endometabolites and bacterial transcripts. Of 31 phytoplankton endometabolites identified and classified into temporal abundance patterns, eight could be matched to patterns of bacterial transcripts mediating their uptake and catabolism. A model simulating the coupled endometabolite-transcription relationships hypothesized that one category of outcomes required an increase in phytoplankton metabolite synthesis in response to the presence of the bacterium. An experimental test of this hypothesis confirmed higher endometabolome accumulation in the presence of bacteria for all five compounds assigned to this category – leucine, glycerol-3-phosphate, glucose, and the organic sulfur compounds dihydroxypropanesulfonate and dimethylsulfoniopropionate. Partitioning of photosynthate into rapidly-cycling dissolved organic molecules at the expense of phytoplankton biomass production has implications for carbon sequestration in the deep ocean. That heterotrophic bacteria can impact this partitioning suggests a previously unrecognized influence on the ocean’s carbon reservoirs.

**Significance Statement:** Microbes living in the surface ocean are critical players in the global carbon cycle, carrying out a particularly key role in the flux of carbon between the ocean and atmosphere. The release of metabolites by marine phytoplankton and their uptake by heterotrophic bacteria is one of the major routes of microbial carbon turnover. Yet the identity of these metabolites, their concentration in seawater, and the factors that affect their synthesis and release are poorly known. Here we provide experimental evidence that marine heterotrophic bacteria can affect phytoplankton production and extracellular release of metabolites. This microbial interaction has relevance for the partitioning of photosynthate between dissolved and particulate carbon reservoirs in the ocean, an important factor in oceanic carbon sequestration.

## Introduction

Photoautotroph-heterotroph metabolite transfer in the surface ocean is a key process in global carbon cycling through which up to half of fixed carbon is transferred to bacteria in the form of labile dissolved compounds^1^. Phytoplankton synthesis and release of metabolites exhibit diel cycles, synchronized with the availability of light energy^2–4^. The observation of similar diel activity cycles in co-occurring heterotrophic bacteria suggests a tight temporal linkage controlled by the timing of phytoplankton extracellular release^5,6^. Though the importance of the trophic link between marine phytoplankton and bacteria in the global carbon cycle has long been recognized^7–9^, identifying the metabolites responsible and measuring their flux is challenging. These compounds have short turnover times in seawater due to rapid uptake by bacteria, have shared physical properties with salt, and are maintained at low, typically nmol L^-1^ to pmol L^-1^, concentrations^10,11^.

Intracellular phytoplankton metabolite pools (endometabolites) are the presumptive substrates for heterotrophic bacteria, yet how faithfully phytoplankton internal concentrations predict exometabolite availability depends on the mechanism of release, of which several have been recognized^12,13^. In the simplest mechanism, differences in metabolite concentration between phytoplankton intracellular pools and ambient seawater can drive diffusion^14^ (i.e., passive diffusion mechanism), in which case substrate supply to heterotrophic bacteria is largely controlled by endometabolite concentrations. Alternatively, active excretion of metabolites to maintain cellular balance can occur by overflow pathways^15^, for example to manage redox state or products of photorespiration (i.e., physiological balance mechanism). Finally, metabolites may be synthesized and excreted in response to associated microbes, for example to sustain mutualisms or mount defenses^16,17^ (i.e., interaction response mechanism).

Here we determined the correspondence between phytoplankton intracellular pools and heterotrophic bacterial substrate availability by examining diel patterns of endometabolomes and transcriptomes. A synthetic community was established in which marine diatom *Thalassiosira pseudonana* CCMP1335^18^ was the only source of substrates to bacterium *Ruegeria pomeroyi* DSS-3^19^. As diatoms contribute up to 40% of primary production in the surface ocean^20^ and *R. pomeroyi* belongs to a taxon that dominates diatom bloom communities^21,22^, this simple community represents a key phytoplankton-bacteria link in the surface ocean. Over two day-night cycles, we contemporaneously assayed phytoplankton endometabolite pools by nuclear magnetic resonance (NMR) spectroscopy and bacterial metabolite consumption using transcriptome proxies and assessed links between the two. Transcript abundance was analyzed as the number of mRNA molecules per bacterial cell, enabled by the use of internal mRNA standards; this approach yields the absolute number of transcripts harbored by a cell for a given gene, matching absolute quantitation in the metabolite data and eliminating ambiguities inherent in proportional expression data^23,24^. The quantitative chemical-biological analytical framework applied to this synthetic community enabled us to assess mechanisms underlying temporal links between microbial autotrophs and heterotroph in the production and consumption of labile metabolites.

## Results and Discussion

*T. pseudonana* cultures were grown axenically under naturally oscillating light intensity during a 16 h:8 h light:dark cycle, with maximum at noon. After 6 d, *R. pomeroyi* was inoculated into the cultures, and a 2-day pre-incubation followed to allow the bacteria to assimilate labile metabolites that accumulated during the axenic phase. Beginning on day 8, samples were collected every 6 h for the next 48 h at timepoints corresponding to midnight, mid-morning, noon, and mid-afternoon.

### Diatom metabolome composition

NMR characterization of the diatom endometabolome during the 48 h sampling window revealed 282 major peaks absent in the blank spectrum. Annotation by comparison to metabolomic databases and chemical standards suggested by previous studies^25,26^ resulted in 31 compounds (156 peaks) identified with high confidence (Table 1; see Table S1 and Fig. S1 for detailed annotation and confidence level information). The number of diatom cells increased ~2-fold over the sampling window, from 0.87 to 1.9 × 10^5^ cells mL^-1^ (Fig. 1a); metabolite data were normalized to cell number at the time of sampling.

**Figure 1.**
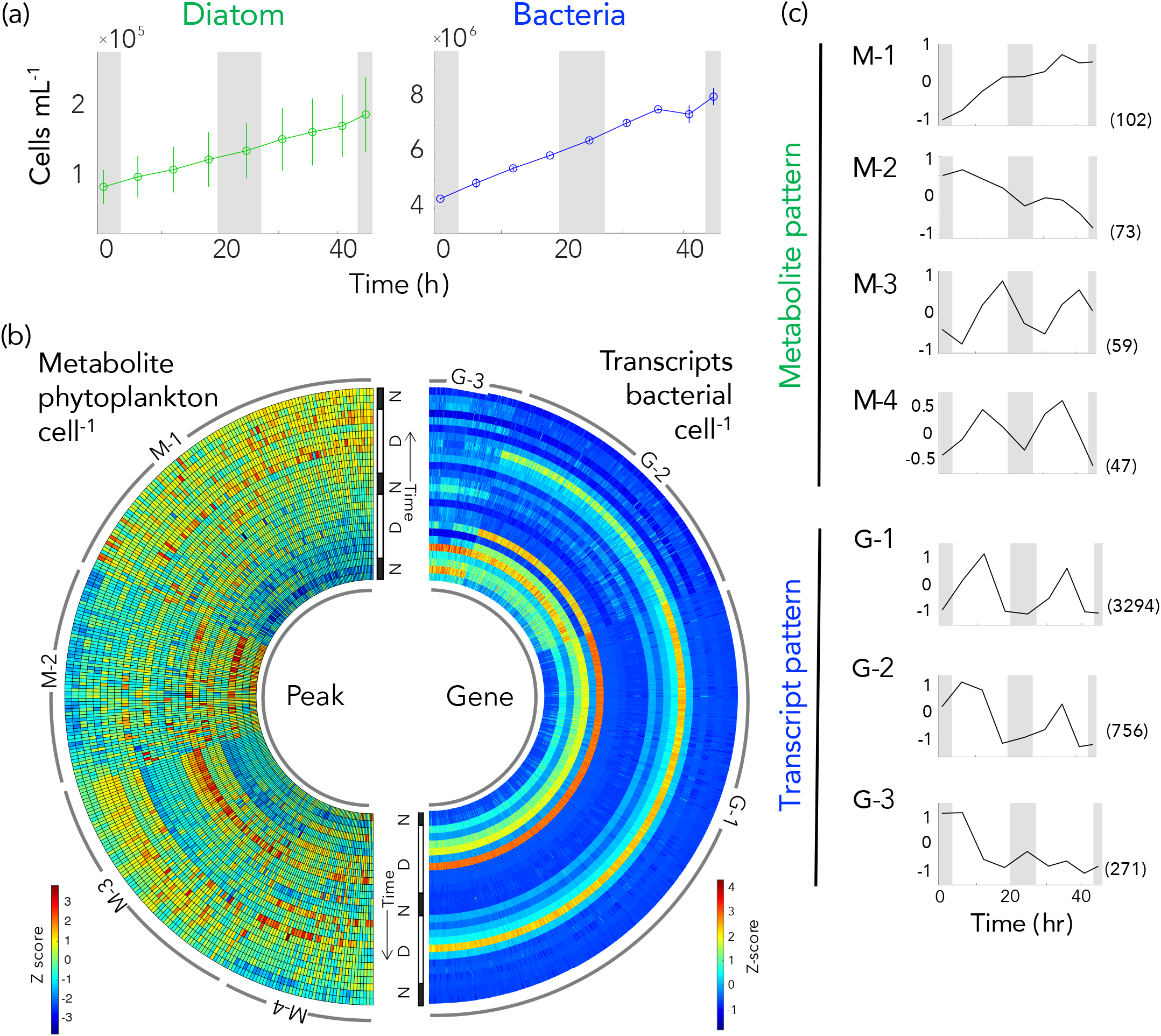
(a) Cell numbers of co-cultured diatoms and bacteria. (b) Temporal variations in metabolite concentration per diatom cell (left) and transcripts per bacterial cell (right) for genes differentially expressed between noon and night (≥2 fold-change and DESeq2 adjusted-p ≤ 0.05). Values were converted to Z-scores and data from each of the three biological replicates are shown. (c) Temporal patterns identified for metabolites (M-1 through M-4) and gene transcription (G-1 through G-3). The number of metabolite peaks or genes in each cluster is given in parentheses. Grey shading in panels a and c indicates night.

**Table 1.**
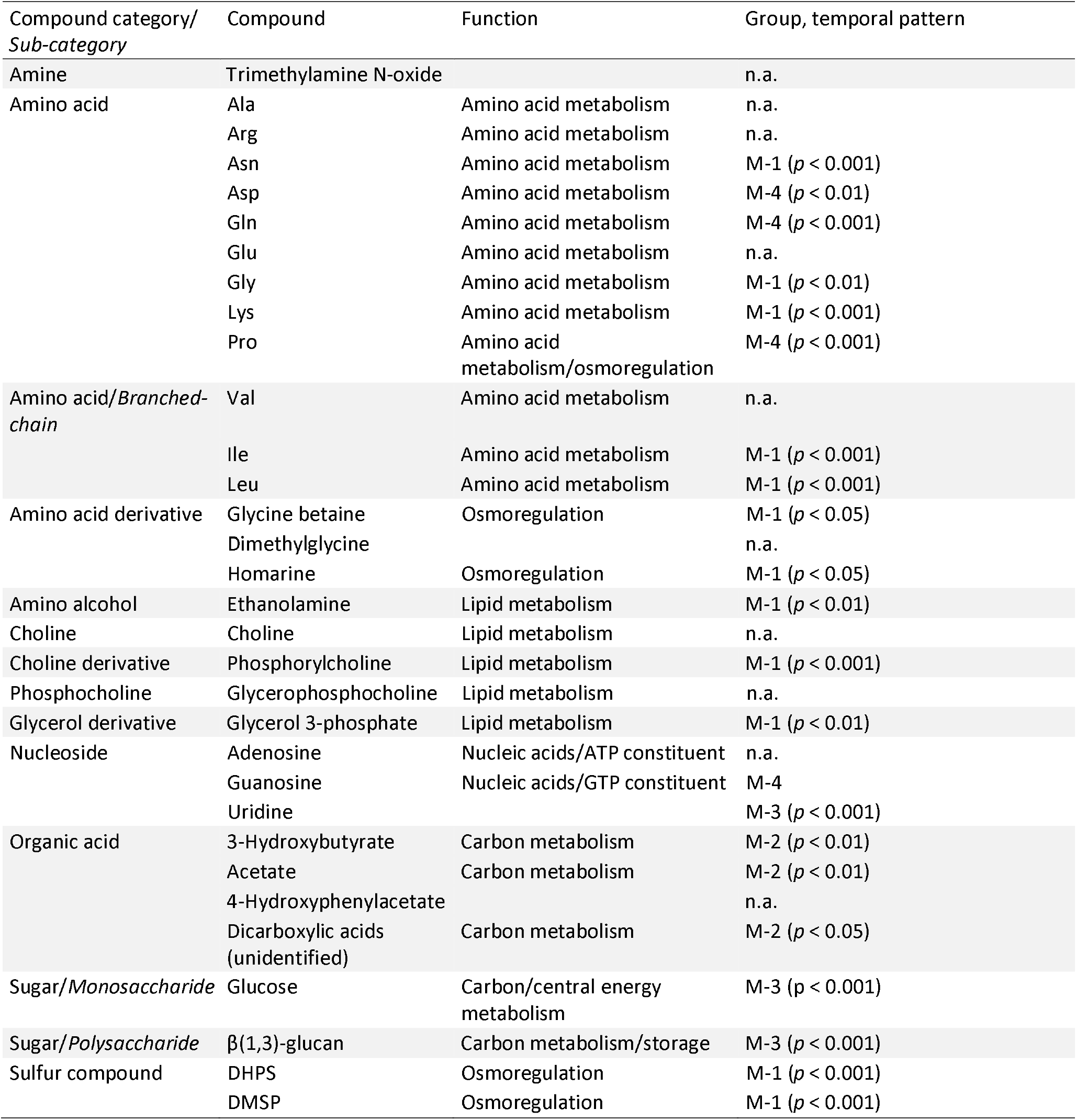
Diatom endometabolites assigned with high confidence in the diel experiment. For detailed information for compound identification and confidence level information, see Table S1 and Fig. S1. Group assignments correspond to those in Figure 1. Statistical significance for temporal patterns is based on linear regression analysis for increasing or decreasing patterns, and RAIN for diel cycles. Temporal patterns: M1 = increase, M2 = decrease, M3 = diel with a peak at mid-afternoon, M4 = diel with a peak at noon. n.a., not applicable (membership value of <0.5, see text for the detail).

| Compound category/<br>Sub-category | Compound | Function | Group, temporal pattern |
| --- | --- | --- | --- |
| Amine | Trimethylamine N-oxide |  | n.a. |
| Amino acid | Ala | Amino acid metabolism | n.a. |
|  | Arg | Amino acid metabolism | n.a. |
| | Asn | Amino acid metabolism | M-1 ($p < 0.001$) |
| | Asp | Amino acid metabolism | M-4 ($p < 0.01$) |
| | Gln | Amino acid metabolism | M-4 ($p < 0.001$) |
|  | Glu | Amino acid metabolism | n.a. |
| | Gly | Amino acid metabolism | M-1 ($p < 0.01$) |
| | Lys | Amino acid metabolism | M-1 ($p < 0.001$) |
| | Pro | Amino acid metabolism/osmoregulation | M-4 ($p < 0.001$) |
| Amino acid/Branched-chain | Val | Amino acid metabolism | n.a. |
| | Ile | Amino acid metabolism | M-1 ($p < 0.001$) |
| | Leu | Amino acid metabolism | M-1 ($p < 0.001$) |
| Amino acid derivative | Glycine betaine | Osmoregulation | M-1 ($p < 0.05$) |
|  | Dimethylglycine |  | n.a. |
| | Homarine | Osmoregulation | M-1 ($p < 0.05$) |
| Amino alcohol | Ethanolamine | Lipid metabolism | M-1 ($p < 0.01$) |
| Choline | Choline | Lipid metabolism | n.a. |
| Choline derivative | Phosphorylcholine | Lipid metabolism | M-1 ($p < 0.001$) |
| Phosphocholine | Glycerophosphocholine | Lipid metabolism | n.a. |
| Glycerol derivative | Glycerol 3-phosphate | Lipid metabolism | M-1 ($p < 0.01$) |
| Nucleoside | Adenosine | Nucleic acids/ATP constituent | n.a. |
|  | Guanosine | Nucleic acids/GTP constituent | M-4 |
| | Uridine | | M-3 ($p < 0.001$) |
| Organic acid | 3-Hydroxybutyrate | Carbon metabolism | M-2 ($p < 0.01$) |
| | Acetate | Carbon metabolism | M-2 ($p < 0.01$) |
|  | 4-Hydroxyphenylacetate |  | n.a. |
| | Dicarboxylic acids (unidentified) | Carbon metabolism | M-2 ($p < 0.05$) |
| Sugar/Monosaccharide | Glucose | Carbon/central energy metabolism | M-3 ($p < 0.001$) |
| Sugar/Polysaccharide | $\beta$(1,3)-glucan | Carbon metabolism/storage | M-3 ($p < 0.001$) |
| Sulfur compound | DHPS | Osmoregulation | M-1 ($p < 0.001$) |
| | DMSP | Osmoregulation | M-1 ($p < 0.001$) |

To group metabolite peaks that behaved similarly over the diel cycles, cell-normalized absolute abundance data were clustered by variance-sensitive clustering^27^ which identified four patterns (Fig. 1b and 1c; Table 1). Group M-1 consisted of metabolites for which monotonic increases in intensity dominated the 48 h sampling window. Twelve compounds annotated with high confidence from this cluster included amino acids (asparagine, glycine, isoleucine, leucine, and lysine), amino acid derivatives (glycine betaine and homarine), an amino alcohol (ethanolamine), a choline derivative (phosphorylcholine), a glycerol derivative (glycerol-3-phosphate), and the sulfur-containing compounds dihydroxypropanesulfonate (DHPS) and dimethylsulfoniopropionate (DMSP) (Table 1, Fig. S2). Metabolite group M-2 was characterized by peaks for which decreases in concentrations over time was the dominant pattern, and included two organic acids (3-hydroxybutyrate, acetate) and one unidentified organic acid. The two other metabolite clusters exhibited diel concentration patterns that peaked in the light and declined in the dark (Table 1, Fig. S2). Group M-3 peaks reached their maximum intensities at mid-afternoon (RAIN, *p* ≤ 0.001) and contained high-confidence annotations of the nucleoside uridine and the carbohydrates glucose and β-1,3-glucan, the latter a subunit of the major diatom polysaccharide laminarin^28^. Group M-4 peaks exhibited diel patterns with maximum intensities at mid-morning or noon (RAIN, *p* ≤ 0.01) and included high confidence annotations for the amino acids aspartate, glutamine, and proline. Thus four distinct temporal patterns of endometabolite concentrations were observed for *T. pseudonana* cells co-growing with a heterotrophic bacterium under a light regime mimicking that of the surface ocean (Fig. 1b).

### Bacterial transcription patterns

We next examined concurrent bacterial transcript inventories indicative of metabolite consumption, normalized to cell counts at the time of sampling (Fig. 1a). The total number of transcripts cell^-1^ varied significantly over the diel cycle (ANOVA; n = 26, p < 0.01), with ~2.5-fold more mRNAs in the mid-morning and noon cells (95 ± 49 and 114 ± 53 mRNAs cell^-1^) relative to mid-afternoon and night (42 ± 11 and 58 ± 25 mRNAs cell^-1^). Correspondingly, the majority of genes had higher transcripts per cell at mid-morning and noon relative to mid-afternoon and night (Fig. S3). This transcript inventory is low compared to exponentially growing *Escherichia coli* (1,350 mRNAs cell^-1^; ref^29^) but comparable to previous measures for marine bacteria in ocean environments.

To identify genes that behaved similarly over time, the per cell transcript inventories for each of the 4,278 protein-encoding genes in the *R. pomeroyi* genome were clustered by variance-sensitive clustering (Fig. 1b and c). Among the genes encoding substrate transporters, the majority (87%) were classified into Group G-1 (3,294 total genes, 539 transporter genes), for which the transcription pattern was a diel cycle with a maximum value at noon (Fig. 2). G-1 transporters showing the largest diel shifts in expression encoded the uptake of sugars (e.g., ribose), amino acid derivatives (ectoine and 5-hydroxyectoine), amines (trimethylamine, trimethylamine-N-oxide, and spermidine), an organic acid (glycolate), a purine (xanthine), phosphonates, and organic sulfur compounds (DHPS, isethionate, cysteate, *N*-acetyltaurine, choline-O-sulfate, and DMSP); for these compounds, bacteria expressed 13- to 58-fold more transcripts per cell at noon relative to night (mean ratio: 33.5 ± 11.7, *n* = 50) (Fig. 2). Transporters with less extreme diel swings in expression but still biased toward noon encoded uptake of taurine, glucose, and *sn*-glycerol-3-phosphate, with noon-night expression ratios an order of magnitude lower (mean ratio: 3.3 ± 1.3, *n* = 11). For these, the dampened diel expression dynamics were due to high night transcript inventories rather than low noon inventories (Fig. 2), suggesting their targets were among the more available substrates at night. Group G-2, for which the temporal transcription pattern was similar to G-1 but with higher values at the first night and mid-morning time points, contained 11% of transporter genes (756 total genes, 68 transporter genes) (Fig. 1c). Group G-3, for which diel patterns were not dominant but, similar to G-2, the first night and mid-morning values were high (Fig. 1c), contained 2% of transporter genes (271 total genes, 10 transporter genes). Putrescine, glycine betaine/proline, and choline transporter proteins were classified in G-2 or G-3. Higher transcript inventories at initial time points could reflect incomplete bacterial drawdown of an accumulated metabolite during the pre-incubation.

**Figure 2.**
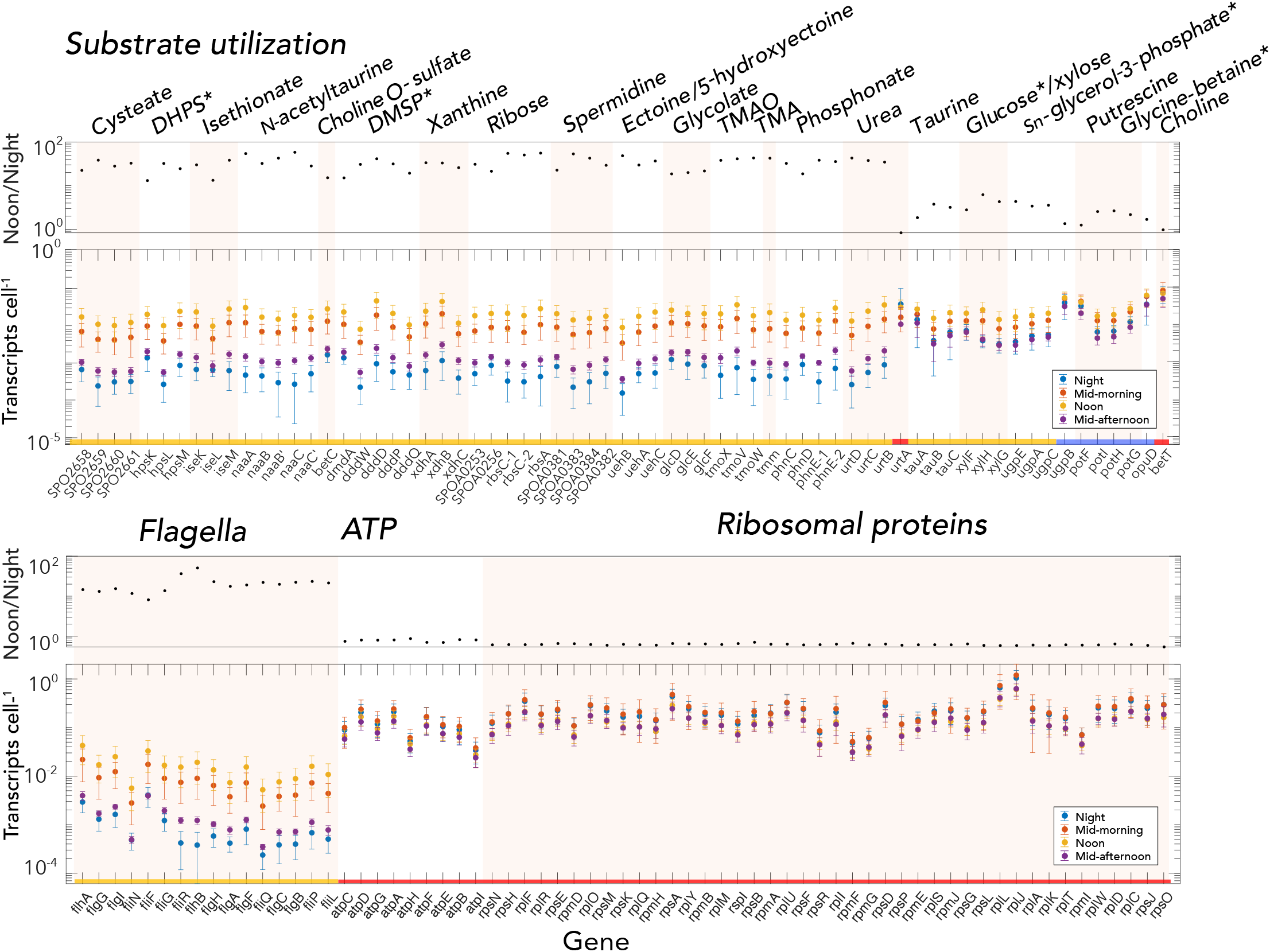
Expression levels of representative *R. pomeroyi* genes encoding transporters or diagnostic catabolic genes (top) and flagella, ATPases, and ribosomal proteins (bottom). For each panel, the top plot shows noon to night ratios (black circles), and the bottom plot shows average transcripts cell^-1^ at night, mid-morning, noon, and mid-afternoon. Error bars indicate standard deviations. Categories of transcription temporal patterns (G-1, gold; G-2, blue, G-3, red) are indicated along the x-axis. Asterisks indicate transporters whose target substrate matches an endometabolite identified with high confidence.

Bacterial transporter expression was also calculated as a percent of the total transcriptome (Fig. S4), the prevailing analysis approach for RNAseq data^30^ when internal standards are not available. Relative investment calculations categorized 51% of transporter genes as significantly enriched in the noon transcriptome relative to night, compared to 81% significantly higher per cell transcript inventories at noon relative to night for the internal standard-based approach (Fig. S4). These analyses emphasize, on the one hand, the bacterium’s investment in expression of a transporter relative to other cellular functions, and on the other, the actual number of templates available for synthesizing transporter proteins.

### Diel oscillations in photosynthesis parameters

We noticed that *R. pomeroyi* produced many-fold fewer transcripts for substrate acquisition in the mid-afternoon compared to mid-morning (Fig. 2), despite the fact that illumination was identical. Indeed, >75% of the bacterium’s transporter genes had transcript inventories that were statistically indistinguishable between mid-afternoon and night (Fig. S3). Diel oscillations in the relationship between carbon fixation rate and irradiance have been broadly documented for marine phytoplankton in laboratory and field studies, characterized by pre-noon maxima in photosynthesis rates^31,32^. Thus the rapid decrease in expression of most bacterial transporters by mid-afternoon suggests that periodicity in carbon fixation-irradiance relationships are manifested in phytoplankton extracellular release as well. One feature of diel photosynthesis oscillation, the E_k_-dependent variability in photosynthesis parameters, is hypothesized to result from a metabolic shift by phytoplankton from pre-noon synthesis of amino acid and lipids to post-noon synthesis of storage carbohydrates and nucleic acids^33,34^. The *T. pseudonana* endometabolome concentrations were fully consistent with this hypothesis; high confidence metabolites assigned to group M-4 (maximum concentrations at mid-morning or noon) are proline, aspartate, and glutamine, and to group M-3 (maximum concentrations at mid-afternoon) are glucose, β(1,3)-glucan storage molecules, and uridine (Fig. S2 and S5). Previously observed offsets in diel timing of maximum transcription by distinct surface ocean bacterial taxa^5^ could thus reflect changes in the composition of phytoplankton extracellular release.

### Control for direct effects of light

Although no light-sensing proteins have been definitively identified in the *R. pomeroyi* genome, we assessed whether light could be directly responsible for changes in gene expression. The bacterium was inoculated into spent medium from axenic *T. pseudonana* and exposed in triplicate to one of three light levels matching co-culture irradiance at noon, mid-morning/mid-afternoon, and night for 4 h. Only 61 genes in the bacterial culture (1.4% of the *R. pomeroyi* genome) were significantly enriched by one or both light levels (Fig. S6), indicating that diel differential expression in the co-cultures was primarily mediated indirectly through phytoplankton activities. One group of 10 light-enriched genes function in protection against reactive oxygen species (ROS) (Fig. S7, Table S2), which can be formed when light interacts with oxygen or organic compounds^35,37^. A second group of 16 enriched genes function in the uptake and metabolism of phosphate (*pstSCAB, phoU*), and phosphonate (*phnDEC, phnIGHLJN*) (Fig. S7), and were likely under the control of the similarly enriched *phoB* regulatory protein^38,40^. Phosphorus acquisition transcript enrichment was surprising, since phosphorus availability was identical at all light levels, and phosphate concentrations remain non-limiting for many weeks in this synthetic culture system (>10 μmol L^-1^)^26^. This raises the possibility of light-dependent stimulation of phosphorus acquisition by bacteria that compete with phytoplankton for nutrients^41^, consistent with temporal partitioning of nutrient uptake observed in ocean data^42^, and potentially triggered by seawater ROS concentrations.

### Coincidence of diatom metabolite accumulation and bacterial transcription

Eight metabolites that were represented in the diatom endometabolome dataset had genes recognized to mediate their uptake or catabolism in the bacterial transcriptome dataset (Table 2). For four of these (leucine, glycerol-3-phosphate, DHPS, and DMSP), an increasing endometabolome concentration was paired with a diel gene expression pattern (Figs. 3a, S8). One (proline) exhibited a noon peak in both endometabolome concentration and gene expression; two (glucose and uridine) exhibited mid-afternoon peaks in endometabolome concentration that lagged noon peaks in gene expression by 6 h; and one (acetate) exhibited a decreasing endometabolome concentration paired with diel gene expression (Fig. 3a).

**Figure 3.**
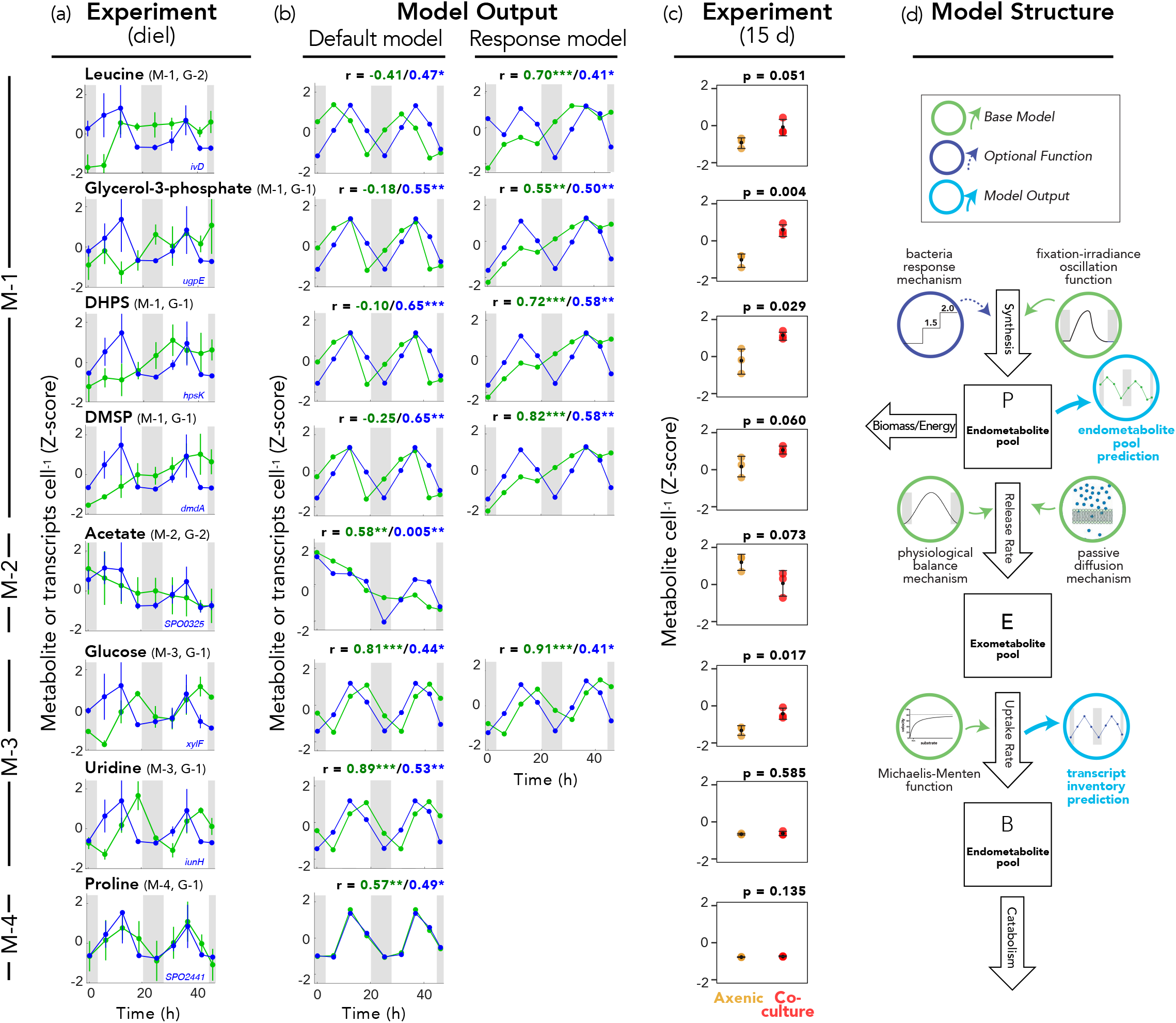
(a) Comparison of temporal patterns for diatom endometabolite concentration (green symbols) and bacterial transcript inventory for a representative gene encoding uptake or catabolism of the same compound (blue symbols); additional relevant genes are shown in Figure S8 (mean + standard deviation, n=3 except for the first night where n=2). (b) Corresponding information from the model output for default (left) and response (right) models. Numbers above the plots indicate r values for Pearson correlations between experimental and model data for metabolite concentrations (green font) and transcript inventories (blue font). *, p≤0.05; **, p≤0.01; ***p≤0.001. (c) Comparison of diatom endometabolite concentrations in axenic culture versus bacterial co-culture (mean + standard deviation, n=3). Numbers above the plots indicate t-test *p* values (n=3). (d) Structure of simulation model.

**Table 2.**
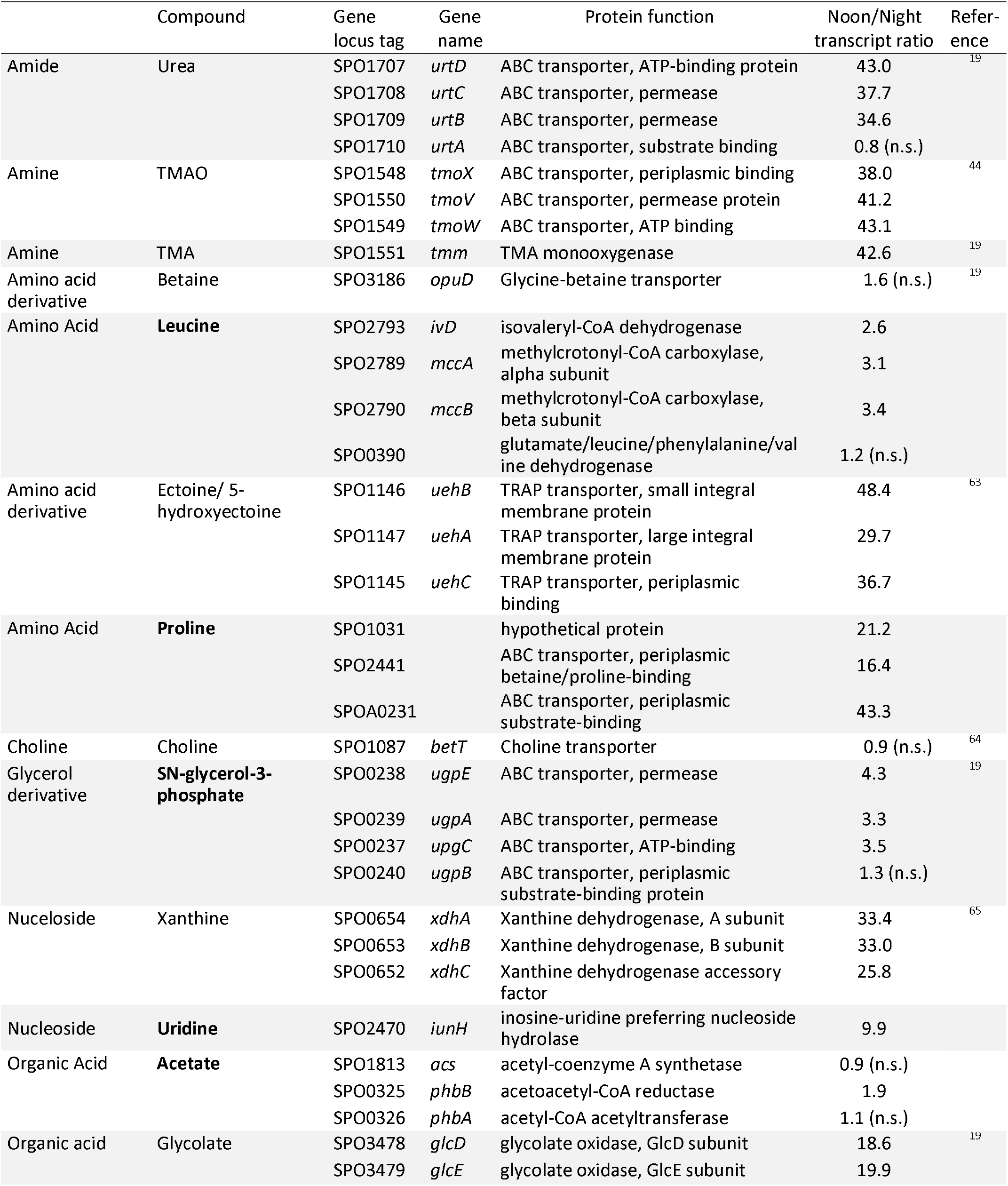
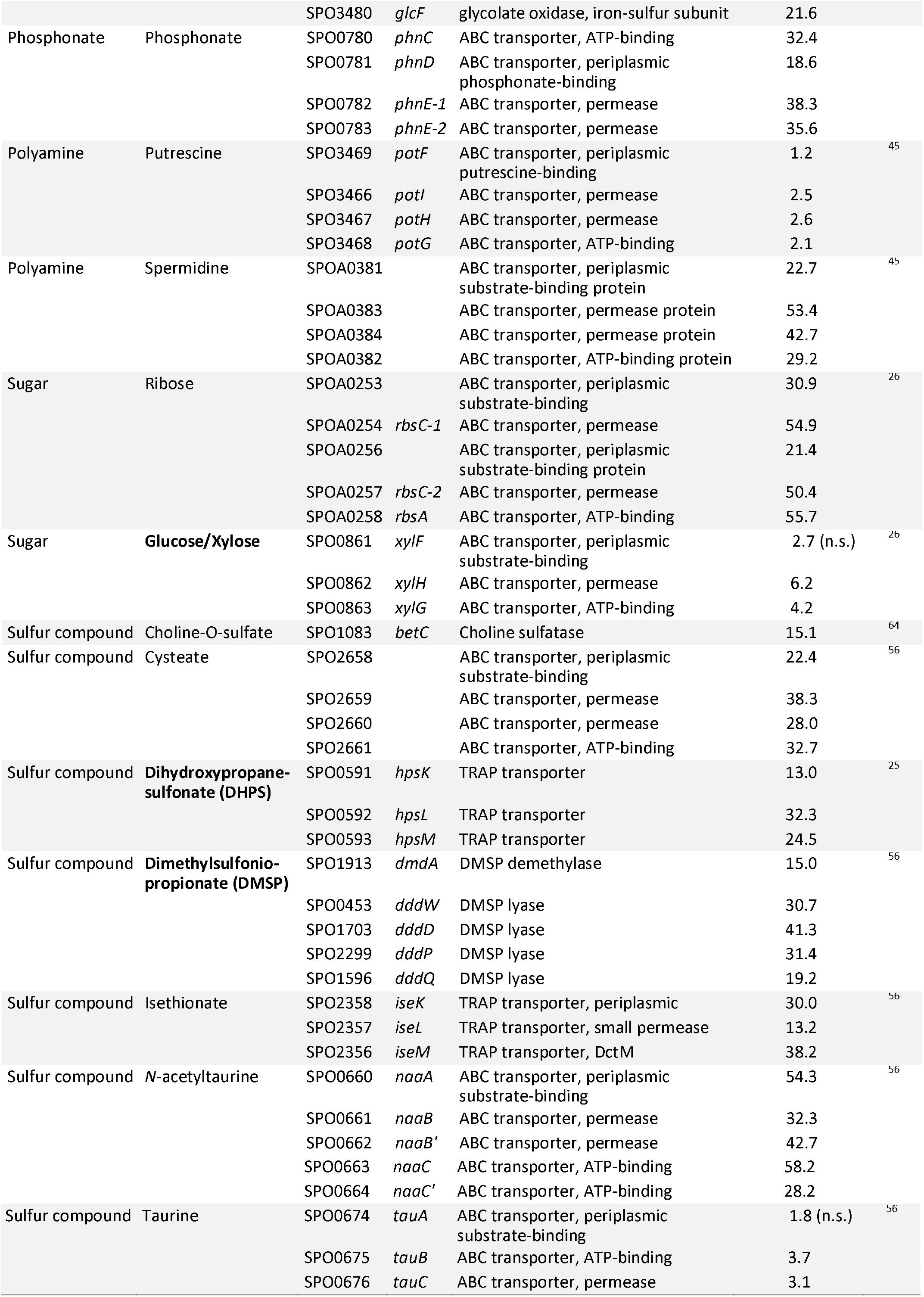
Bacterial noon/night ratios of transcripts cell^-1^ for genes indicative of metabolite consumption. Bold font indicates the compounds appearing in both the endometabolite and bacterial gene expression datasets, n.s., difference not statistically significant (adjusted *p* > 0.05).

We asked whether the observed paired data patterns could occur under a null model of changing phytoplankton exometabolite release following bacterial inoculation; that is, with no mechanism for increased phytoplankton excretion in response to neighboring microbes. A simulation model was used to compute phytoplankton exometabolite release using functions representing two of the proposed mechanisms of extracellular release, passive diffusion and physiological balance, but not the interaction response mechanism (Fig. 3d). To simulate transcription, the model assumed that *R. pomeroyi* transporter systems are regulated by the availability of their substrate, which has been supported in previous studies^43,47^. Thus bacterial transcript inventories were taken as the simulated exometabolite uptake rate, according to Michaelis-Menten kinetics. The model successfully recapitulated three of the four experimental patterns of paired metabolome concentration and transporter expression data (Fig. 3b left; Table S3). The pattern that could not be generated with the base model was that of increasing endometabolome concentrations (M-1) paired with diel gene expression (G-1), which was observed for leucine, glycerol-3-phosphate, DHPS, and DMSP (Fig. 3b left) (Pearson’s *r* = −0.41 to −0.10). However, addition of a phytoplankton interaction response mechanism that increased endometabolite production rate upon bacterial inoculation enabled the model output to mimic the M-1, G-1 pattern (Fig. 3b right). We noticed that output for glucose (M-3, G-1) insufficiently captured the temporal trend of endometabolome concentration (Fig. 3b left; Table S3), and further analysis identified a significant linear increase in glucose concentrations (*p* < 0.001) embedded within a significant diel pattern (RAIN, *p* < 0.01). Implementation of a phytoplankton interaction response also improved simulation of the glucose data (Fig. 3b right).

The simulation modeling generated a hypothesis that diatoms accumulate higher concentrations of certain endometabolites in the presence of heterotrophic bacteria. This was tested using an available independent dataset in which *T. pseudonana* was grown in co-culture with bacteria (*R. pomeroyi* and two other heterotrophic bacteria) for 15 d, after which endometabolites were compared with those in axenic controls. Consistent with model predictions, all five endometabolites that increased in concentration in the diel study also had higher concentrations in co-culture endometabolomes compared to axenic in the 15 d study (Fig. 3c). For the three metabolites that did not increase in concentration during the diel study (acetate, uridine, and proline) there was no difference in endometabolite concentrations in the 15 d study (Fig. 3c; see also Fig. S9 for other compounds). How *T. pseudonana* might detect associated bacteria is not yet known, but could involve signals released from the bacteria or by bacterial alteration of environmental conditions, such as nutrient pools. Previous research uncovered a phytoplankton-bacteria signaling system in which a marine diatom (*Pseudo-nitzchia multiseries*) released tryptophan extracellularly, and a co-cultured marine bacterium (*Sulfitobacter* sp. S11) converted it to the plant hormone indole-3-acetic acid. In our model system, bacterium *R. pomeroyi* (a relative of *Sulfitobocter* sp. S11, both members of the Roseobacter group) maintained 7- to 36-fold higher transcript inventories of IAA synthesis genes at noon relative to night (Fig. 4a). Further, expression of these same IAA genes was positively correlated with diatom biomass when *R. pomeroyi* was introduced at intervals into a natural phytoplankton bloom^48^ (Fig. 4b). These suggest both that IAA may play a role in a *T. pseudonana - R. pomeroyi* interaction, and that IAA signaling may broadly underlie marine diatom-bacteria interactions in the surface ocean^16^. Our observation that concentrations increase only for certain components of the diatom exometabolome is consistent with evolutionary tuning through selection.

**Figure 4.**
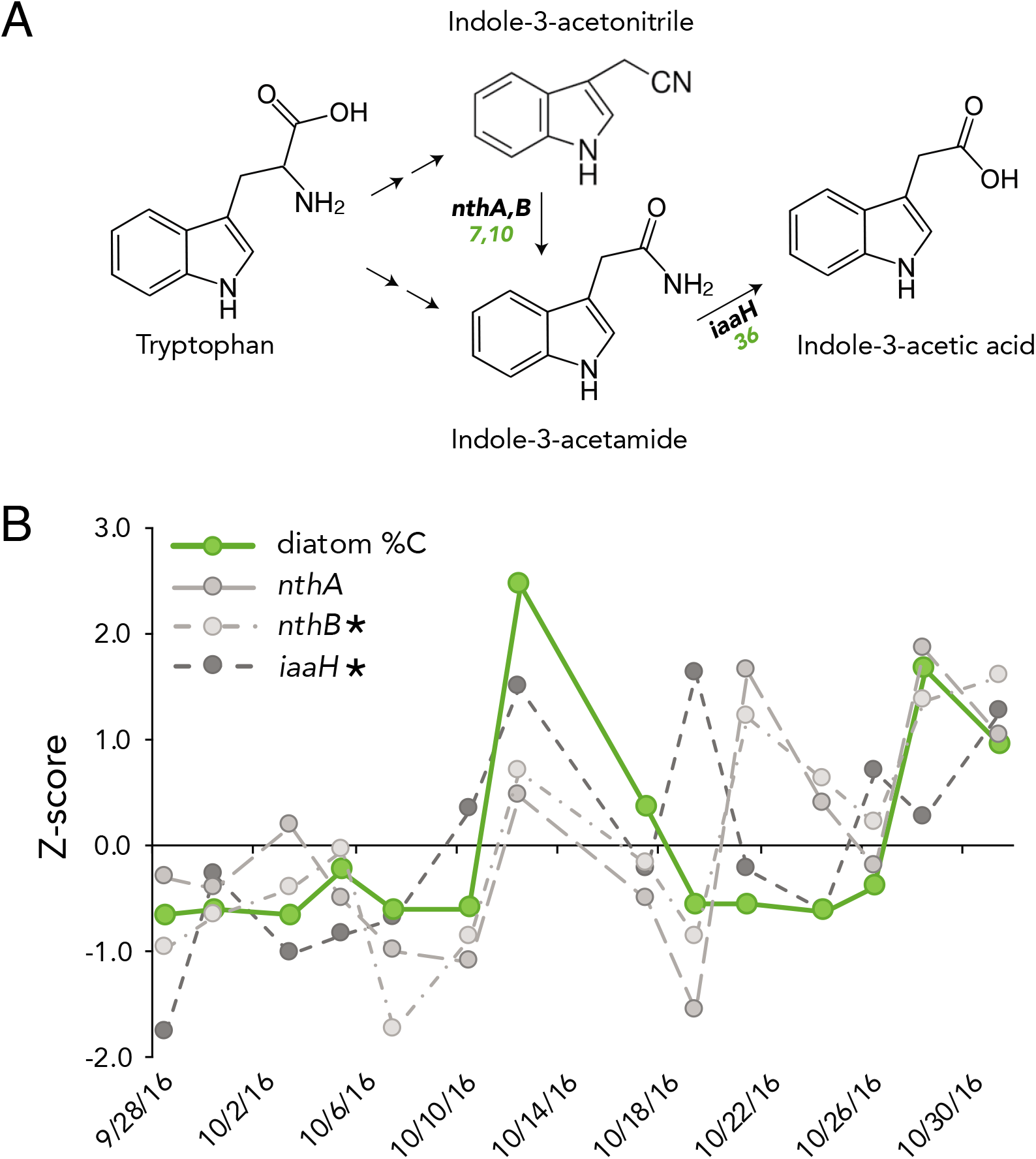
A) Indole-3-acetic acid (IAA) synthesis pathways in *R. pomeroyi. nthA,B*, nitrile hydratase; *iaaH*, IAM hydrolase; green text, noon/night per cell transcript inventories. B) Expression of *R. pomeroyi* IAA synthesis genes (gray symbols) following inoculation into natural phytoplankton bloom communities from Monterey Bay, CA, USA. Bacteria were added at 3-4 d intervals over a 1 mo period, with mRNA retrieved for transcriptome sequencing 90 min after inoculation^48^. The percent of phytoplankton carbon in the bloom community contributed by diatoms (green symbols) was calculated from microscopic cell counts and taxon-specific cell volumes. Data are Z-scores of mean values for three replicates. Asterisks indicate *R. pomeroyi* genes whose expression is positively correlated with diatom % C (Pearson’s R ≥ 0.54; p ≤ 0.05).

A major fraction of the ocean’s annual net primary production is processed through the labile dissolved organic carbon pool, driven by the production, release, and consumption of microbial metabolites^9^. In agreement with previous field observations of coincident diel patterns of phytoplankton and bacterial activity^5,42,49^, we find coupling of phytoplankton endometabolite dynamics with bacterial exometabolite uptake transcription in a model system representative of diatom-dominated surface ocean ecosystems. The quantitative importance of marine phytoplankton-bacteria carbon flux has motivated inquiries into the physical and chemical factors that regulate phytoplankton extracellular release, such as light, temperature, and nutrient limitation^33^. This study suggests that heterotrophic bacteria also influence this process, with implications for ocean carbon sequestration via allocation of photosynthate between dissolved and particulate organic carbon reservoirs.

## Materials and Methods

### Diel experiment

An axenic strain of marine diatom *Thalassiosira pseudonana* CCMP1335 was cultured at 18°C in three replicate 15-L polycarbonate bottles containing 10 L of LI medium^50^ in which NaH^13^CO_3_ (Cambridge Isotope Laboratories, CLM-441) was used as the source of inorganic carbon. The light cycle consisted of 16 h light, during which light intensity varied gradually between 0 and 150 μmol photon m^-2^ s^-1^ with a maximum intensity at noon, followed by 8 h of dark. Bacterial strain *Ruegeria pomeroyi* DSS-3 was grown at 30°C on ½ YTSS agar and transferred to ½ YTSS liquid medium for overnight growth. Axenic *T. pseudonana* cultures grown for 6 days were inoculated with bacterial cells washed in L1 medium three times (final concentration, 10^6^ bacterial cells mL^-1^). Cocultures were pre-incubated for two days to allow time for accumulated labile phytoplankton metabolites to be consumed by the bacteria and thus emphasize synchronized production and consumption dynamics during diel cycles. After the pre-incubation period, samples were collected every 6 h over the next 48 h for bacterial mRNA sequencing, phytoplankton and bacterial cell counts, and phytoplankton endometabolome analysis.

### Direct light effects experiment

*T. pseudonana* CCMP1335 was axenically cultured in 10 L of L1 medium in a 15-L polycarbonate bottle with incubation conditions as described above except that an intensity of 150 μmol photon m^-2^ s^-1^ was used throughout the light period. After one week, the diatom cultures were sequentially filtered through GF/F filters (Whatman) and 0.2-μm-pore-size PCTE membrane filters (Poretics), and the cell-free filtrate was used as the substrate for a bacterial monoculture experiment. *R. pomeroyi* DSS-3 cells were prepared and added to the filtrate as described above. Cells were incubated for 4 h at 18°C under light intensities of 150 (100% treatment), 75 (50% treatment), or 0 μmol photon m^-2^ s^-1^ (0% treatment), corresponding to light levels at noon, mid-morning and mid-afternoon, and night in the diel experiment, with three replicates of each treatment. A minor temperature increase of O.5°C occurred in the 100% treatment relative to 50% and 0% treatments. After 4 h, samples for bacterial RNA analysis and cell counts were collected.

### Diatom endometabolome analysis

Diatom cells were collected by filtering 500 mL of culture onto 2.0-μm-pore-size PCTE membrane filters (MilliporeSigma Isopore) and stored at −80°C until processing. Endometabolites were extracted by sonication in ultra-pure water (Millipore), concentrated by freeze-drying, and dissolved in 600 μL of sodium phosphate butter (pH 7.4) with an internal standard of 2,2-dimethyl-2-silapentane-5-sulfonate-d_6_ (1 mmol L^-1^)^51^. Metabolites were analyzed by nuclear magnetic resonance (NMR) spectroscopy using a Bruker AVANCE III 800 MHz 5 mm TCI cryoprobe, 800 MHz 1.7 mm TCI cryoprobe, and 600 MHz 5 mm TXI probe. Pulse programs of ^1^H-^13^C heteronuclear single quantum correlation (HSQC; Bruker program hsqcetgpprsisp2.2), ^1^H- ^13^C HSQC-total correlation spectroscopy (HSQC-TOCSY; hsqcdietgpsisp.2), and ^1^H-^13^C heteronuclear multiple bound correlation (HMBC; hmbcetgpl2nd) were used. Data were processed using TopSpin 4.0.3 (Bruker), and peak intensity was extracted using rNMR 1.1.9^52^. Metabolites were annotated based on chemical shift (HSQC) and coupling information (HSQC-TOCSY and HMBC). HMDB^53^ and BMRB^54^ were used as reference databases, and additionally CSDB^55^ for polysaccharides. Three compounds of interest which are not in these databases were annotated either by obtaining original spectra from chemical standards (DHPS and DMSP)^56^ or based on literature values^57^. Confidence level of annotation ranging from 1 (lowest) to 5 (highest) was assigned to each metabolite (Table S1) according to Walejko et al^58^ with a slight modification, where 1 = putative compounds with functional group information; 2 = partially matched to HSQC chemical shift information in the databases or literature; 3 = matched to HSQC chemical shift; 4 = matched to HSQC chemical shift and validated by HSQC-TOCSY or HMBC; 5 = validated by original spectra from chemical standards. Detailed parameter settings are presented in Table S4, with additional information in Metabolomics Workbench (ID PR001019, dx.doi.org/10.21228/M80408). Temporal variations in metabolites were analyzed by extracting peaks behaving similarly during the incubation period using variance-sensitive clustering^27^ after normalization by the internal standard and cell counts, and scaling to Z-scores. Background signals originating from filters and solvent were also corrected. The optimal cluster number was selected based on minimum centroid distance and Xie-Beni index, and only membership values of <0.5 were accepted^27^. Periodicity of the temporal patterns for compounds was analyzed using a rhythmicity analysis package RAIN (1.18.0)^59^ in R software (version 3.6.1). Heatmaps were created using the CirHeatmap function (version 1.7) in MATLAB (Mathworks)^60^.

### mRNA analysis

For the direct light experiment, bacterial cells were collected by filtering 500 mL of culture through 0.2-μm pore-size PES membrane filters (Pall Supor) and immediately freezing the filters in liquid nitrogen. For the diel experiment, samples were pre-filtered through 2.0-μm-pore-size PCTE membrane filters (MilliporeSigma Isopore) to retain diatom cells prior to capturing bacterial cells on 0.2-μm pore-size filters. This process was completed within 15 min of collection. The filters were stored at −80°C until processing. To extract RNA, filters were cut into pieces under sterile conditions and shaken with 0.5 mL of 0.1-mm zirconia/silica beads (BioSpec Products) in 1 mL of Denaturation/Lysis Solution (Life Technologies) for 15 min. RNA was extracted from this lysate using the RNeasy Mini Kit (QIAGEN).

For the diel experiment, we used a phenol-chloroform-isoamyl extraction^26^ after confirming good mRNA recovery from both diatom and bacterial samples. To determine the absolute number of transcripts, two internal mRNA standards (size, 1,000 nt) were added to each sample before extraction and the recovery of the standards was determined following Satinsky et al.^23^. After the extraction, DNA was removed by the Turbo DNA-free Kit (Ambion), rRNA was depleted by Ribo-Zero rRNA Removal Kit (Illumina), and mRNA was purified by RNA Clean & Concentrator-5 (Zymo Research) following the manufacturer’s protocols.

Sequencing was carried out on an Illumina NextSeq 550 (Table S5). rRNA reads were identified by blast+ (NCBI 2.7.1 and 2.8.1 for the direct light experiment and the diel experiment, respectively) against an rRNA sequence database and removed. Remaining reads were mapped to the *R. pomeroyi* genome and quantified using HTSeq^61^. Differentially expressed genes were identified in pairwise comparisons of sampling times (diel experiment) or light levels (direct light experiment) using MATLAB for absolute analysis, and DESeq2^62^ for relative transcript analysis. One of the replicate samples from the initial time point of the experiment was lost; otherwise, n = 3 for all analyses. The number of reads per library averaged 19.2 x 10^6^ (range, 13.3–31.9 x 10^6^) and the percentage of rRNA contamination averaged 17.5% (range, 4.1–38.8%). Recovery of the two internal standards was highly consistent (Pearson’s *r* = 0.96; *p* ≤ 0.001; *n* = 26), accounting for 2.2% of mRNA reads recovered per library. All other statistical analyses were conducted using MATLAB. Fold-change values and temporal pattern categories for all the genes are reported in Table S6.

### Cell counts

A 0.5 mL aliquot of culture was fixed with glutaraldehyde (final concentration, 1%) and kept at −80°C until analysis. Samples were thawed, stained with SYBR Green I (Thermo Fisher Scientific; final concentration, 5 x 10^-4^ of commercial stock), and injected into a CytoFLEX flow cytometer (Beckman Coulter). For phytoplankton counts, samples were analyzed without staining. Data were analyzed using CytExpert (Beckman Coulter), and cell density was calculated based on a separate run of a known concentration of bead standards (Beckman Coulter).

### Model development

The extracellular release model was written in R version 3.6.1 with three state variables, representing the phytoplankton endometabolome (*P*), the exometabolome (*E*), and the bacterial endometabolome (*B*). The time evolution of these pools was calculated at 0.1 h intervals using the following differential equations.

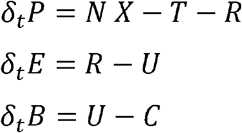

*N* is the metabolite biosynthesis rate, derived from light intensity but allowing for C fixation-irradiance oscillation around the peak in light intensity^32^. *T* is the rate at which endometabolites are allocated for biomass and energy generation by phytoplankton cells, calculated as a constant fraction of *P* at each interval. *R* is release rate of endometabolites from the phytoplankton cell with parameters for both diffusive and physiological balance mechanisms. *U* represents bacterial uptake from the exometabolome following Michaelis-Menten kinetics. *X* is the bacterial response mechanism that increases metabolite biosynthesis rate in the presence of bacteria by 1.5- or 2-fold. C represents catabolism of the metabolite within the bacterial endometabolome, with a constant fraction lost each interval. See Supplemental Methods for information on how variables *N, R, T, U*, and *C* were calculated.

To simulate experimental conditions, *B* and *U* were set to zero for 6 d of ‘axenic growth’ followed by ‘inoculation’ with addition of *B* and *U* functions for the final 4 d of the modeled experiment. Values for *P* and *U* from the final 2 d of model output were used to compare to experimentally measured endometabolome and transcriptome data, respectively.

### Experimental test of model predictions

*T. pseudonana* CCMP1335 was inoculated into L1 medium with NaH^13^CO_3_ labeling as described above. Triplicate samples were inoculated with three heterotrophic bacteria (*Ruegeria pomeroyi* DSS-3, *Stenotrophomonas* sp. SKA-14, and *Polaribacter dokdonensis* MED-152). Another set of triplicate samples was kept axenic (diatom only). The cultures were maintained at 160 μmol photons m^-2^ s^-1^ at 18°C in a 16:8 h light:dark cycle. After 15 d (late stationary phase) diatom cells were filtered from 700 mL of culture, frozen and processed for NMR analysis as described above.

## Data Availability Statement

Data that support the findings of this study have been deposited in NCBI SRA with BioProject accession number PRJNA649292 (sequencing data), and Metabolomics Workbench with Project ID PR001019, dx.doi.org/10.21228/M80408 (metabolome data).

## Acknowledgements

C. Smith and S. Sharma provided sequencing and bioinformatic assistance, J. Gluhska and C. Panagos offered expertise on NMR analysis, M. Landa and B. Nowinski provided valuable comments on experimental design, F. Ferrer-González and J. Schreier assisted with sampling, and the University of Georgia Genomics and Bioinformatics Core (GGBC) provided sequencing services. This work was supported by The Gordon and Betty Moore Foundation (5503), NSF (IOS-1656311), The Simons Foundation (grant 542391 to MAM) within the Principles of Microbial Ecosystems (PriME) Collaborative, JSPS (Research Fellowship for Young Scientists and Grant-in-Aid for JSPS Fellows to MU), and the Swedish Research Council (2018-06571 to MO).

## Author Contributions

MU and MAM conceived of the study, MU and MO collected the data, MU, WS, MO, ASE, and MAM analyzed data, and MU and MAM wrote the paper with input from all authors.

