## Supplemental Material for "Diel Investments in Phytoplankton Metabolite Production Influenced by Associated Heterotrophic Bacteria"

Uchimiya et al.

Supplemental Figures S1-S9

Supplemental Tables S3, S4, and S5

Supplemental Modeling Methods

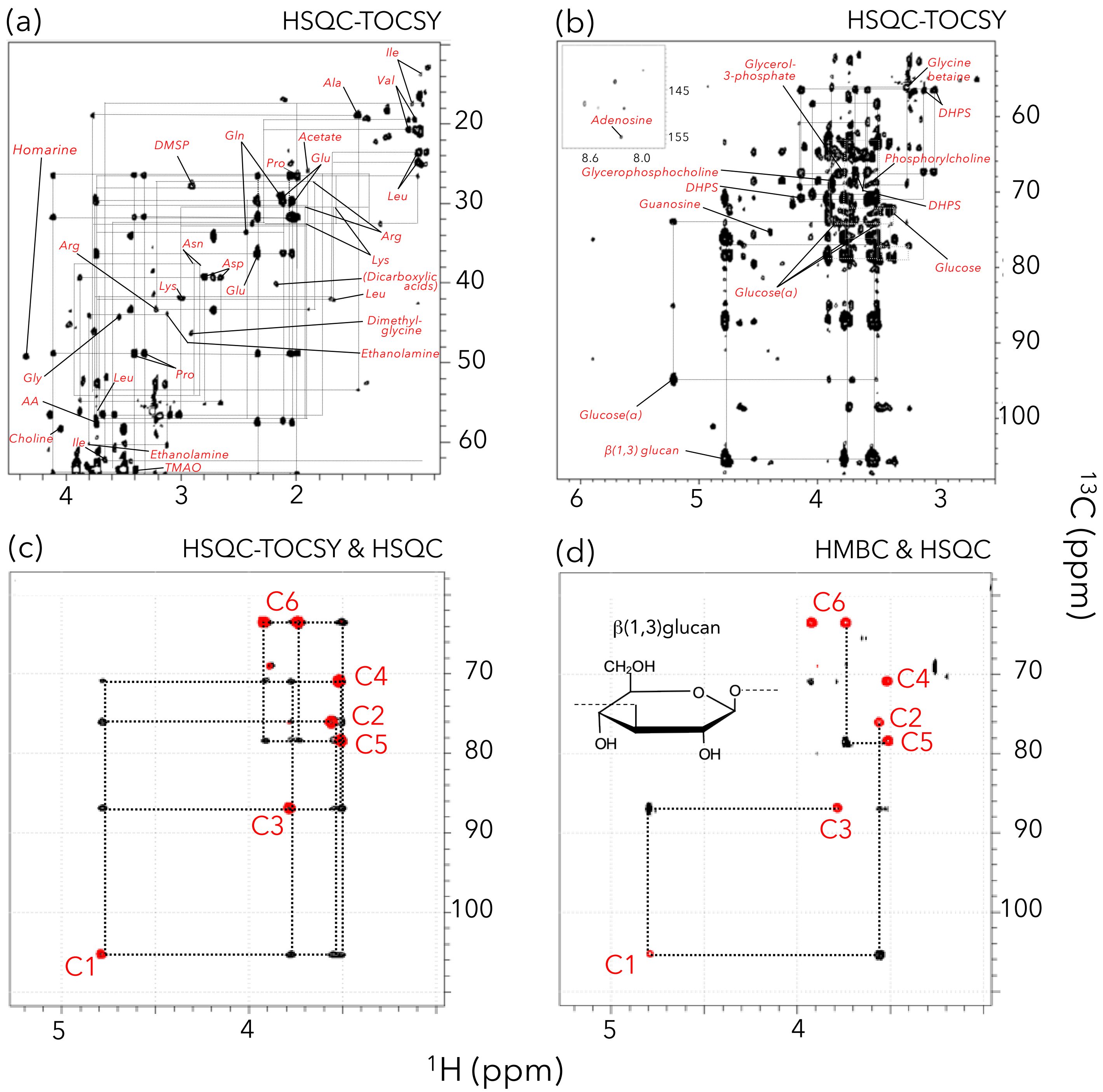


Figure S1. Diatom endometabolome annotation methods. Representative peak(s) for compounds indicated on HSQC-TOCSY spectra (a and b). Additional structural validation (e.g., for polysaccharide β-1,3-glucan) by HSQC-TOCSY (c) and HMBC experiments (d). Peaks from HSQC experiments are overlaid and colored in red. A complete compound list is provided in Table 1, and chemical shift information used for annotation is provided in Table S1. 3-Hydroxybutyrate, 4-hydroxyphenylacetate, and uridine are not visible in a and b due to relatively low intensities. AA, amino acid alpha carbon.

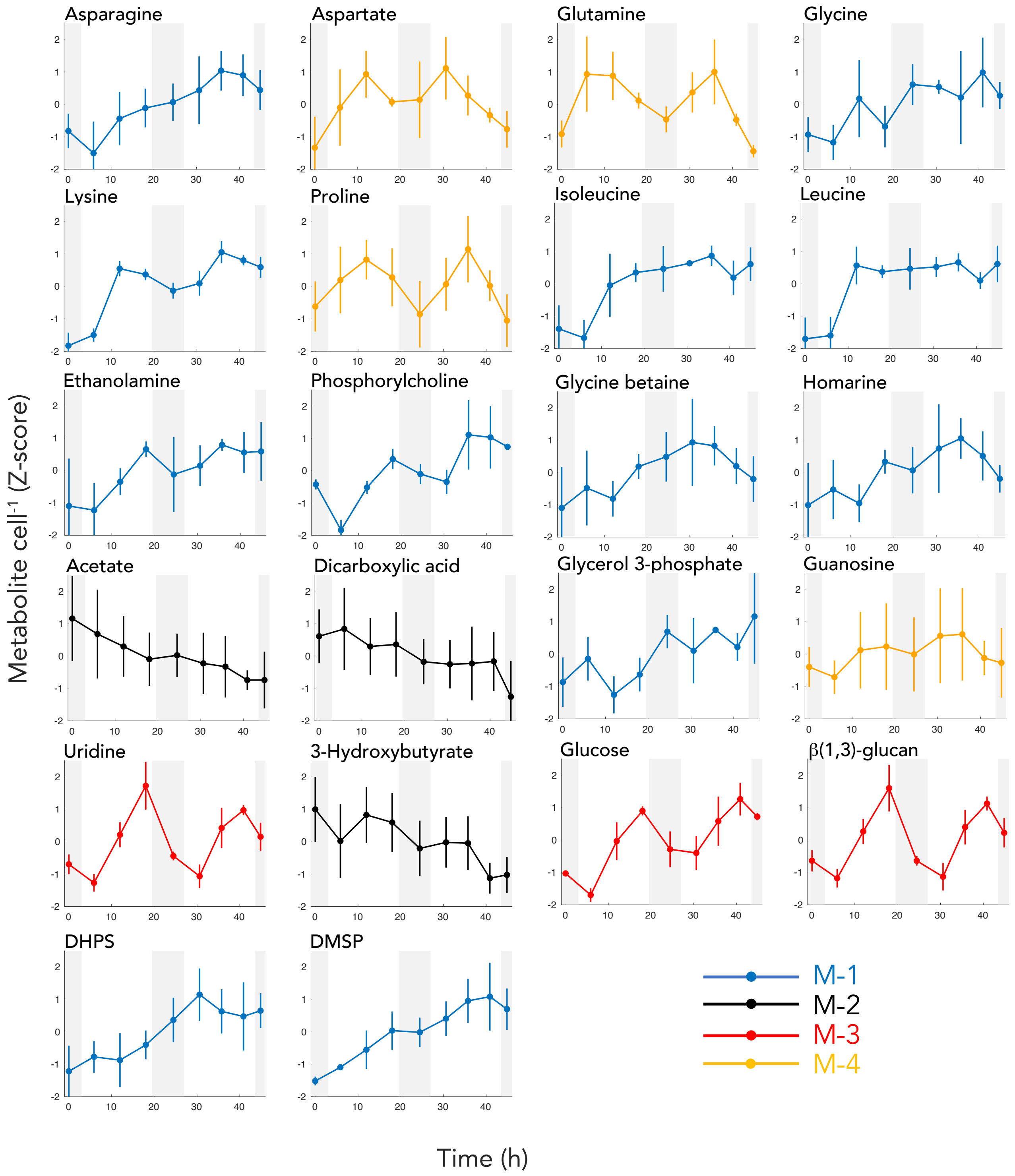


Figure S2. Temporal patterns in 22 diatom endometabolites annotated with high confidence and having significant membership values in a temporal cluster (M-1 through M-4). Metabolite abundance cell$^{-1}$ is shown as Z-scores for one representative non-overlapping NMR peak (see also Table S1). Error bars indicate standard deviation; n=3.

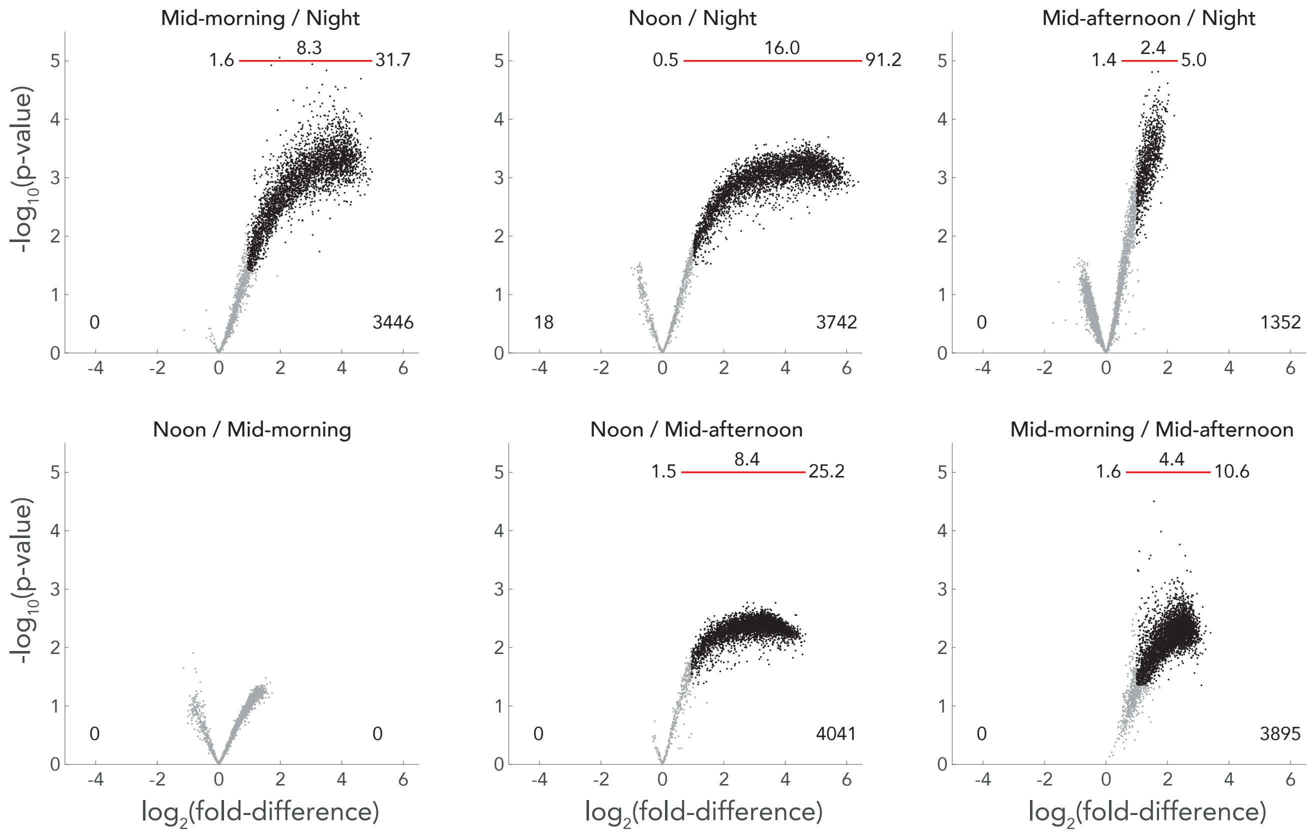


Figure S3. Pair-wise comparison between sample times of bacterial gene expression as transcripts cell$^{-1}$ in the co-culture experiment. Average, minimum- and maximum difference values for differentially expressed genes (black symbols) are shown above the plots. Numbers on the bottom left and right indicate the number of differentially expressed genes (adjusted-p $\leq$0.05 and fold-difference $\geq$2).

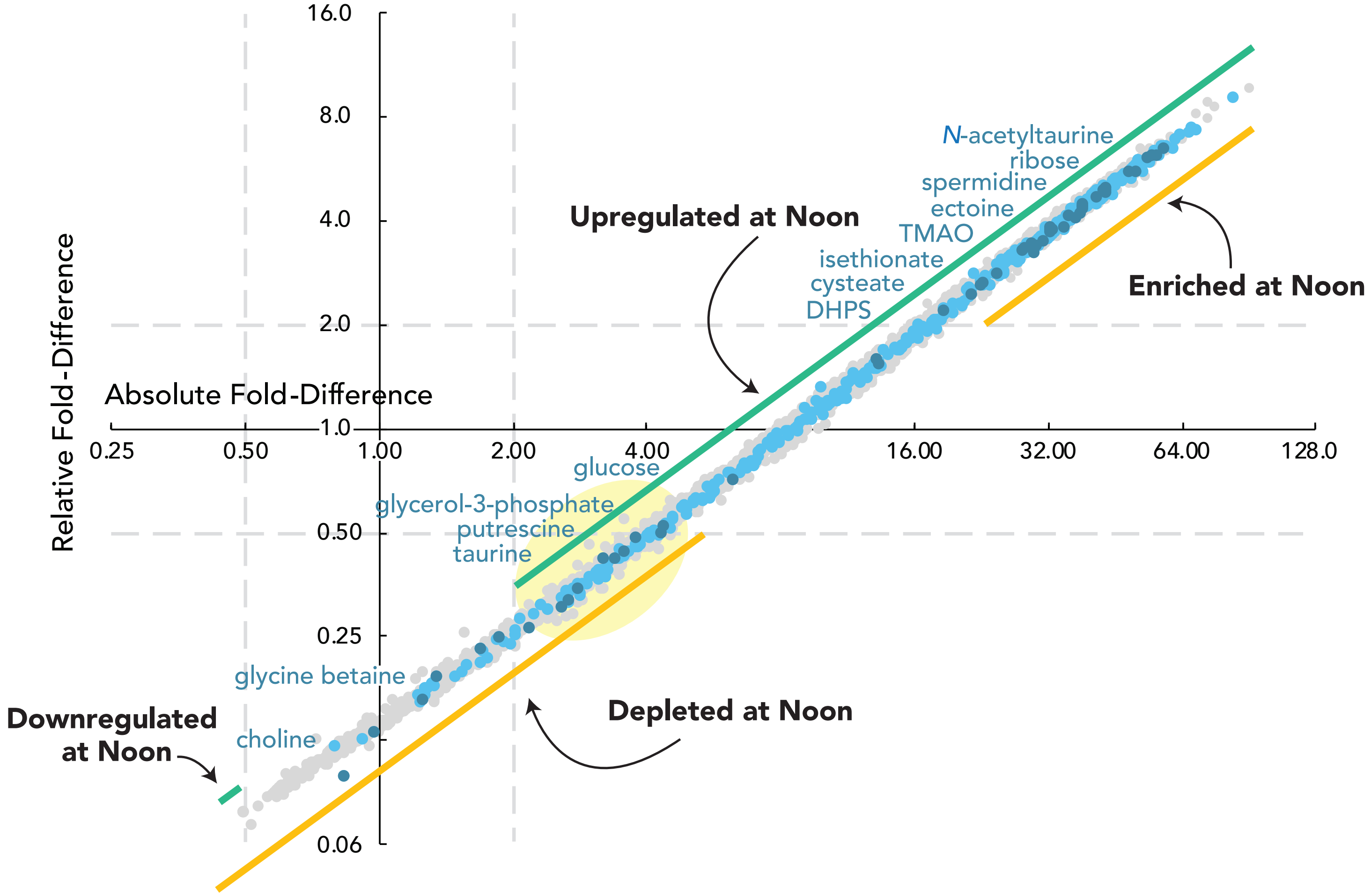


Figure S4. Comparison of fold-difference values for absolute versus relative transcript analysis of noon to night ratios of substrate transporter genes (blue symbols). Absolute analysis (x-axis) represents up- or down-regulation of the number of transcripts per bacterial cell. Relative analysis (y-axis) represents enrichment or depletion as a proportion of the transcriptome. Dark blue symbols indicate the transporter genes in Fig. 2; light blue symbols indicate other transporter genes; grey symbols indicate the remaining *R. pomeroyi* genes. Dashed gray lines mark where fold-difference = 2 on each axis ($log_2$ units). The yellow shading highlights genes with transcript inventories per cell that are significantly higher at noon yet account for a significantly lower proportion of the cells' transcriptome.

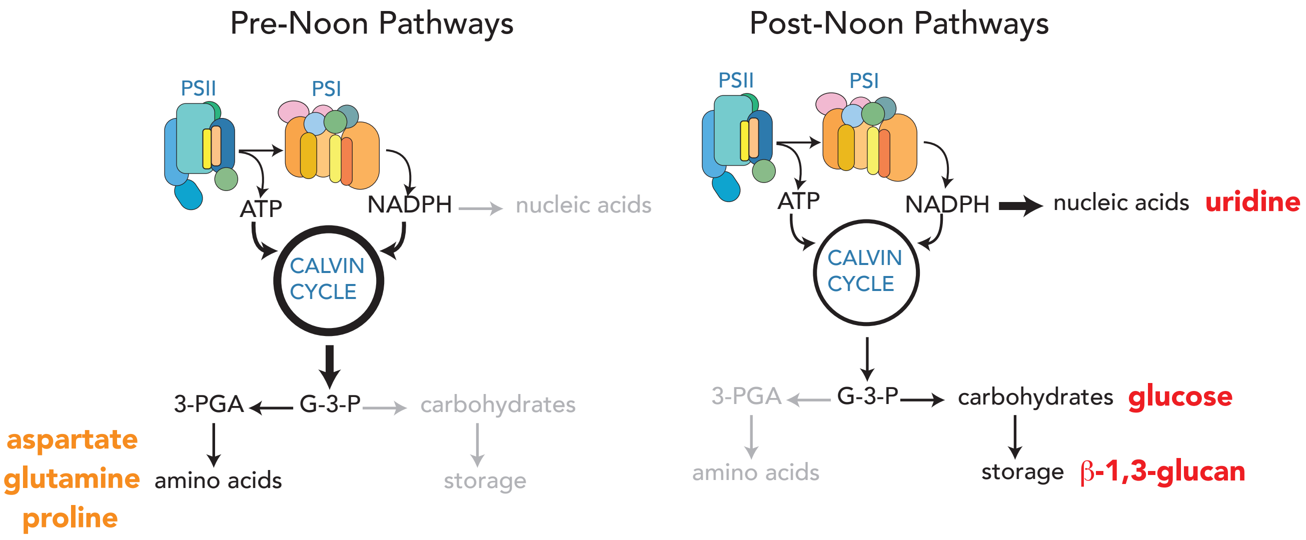


Figure S5. The biochemical pathways hypothesized to dominate marine phytoplankton metabolism pre-noon (left panel) and post-noon (right panel) (Behrenfeld et al. 2008) are good predictors of diel patterns in *T. pseudonana* endometabolite accumulation. Concentrations of aspartate, glutamine, and proline peaked at noon (cluster M-4 metabolites), while concentrations of uridine, glucose, and β-1,3-glucan peaked in mid-afternoon (cluster M-3 metabolites). Temporal endometabolite patterns are shown in Fig. S2. Biochemical pathways are re-drawn from Behrenfeld et al. (2008). G-3-P, glyceraldehyde-3-phosphate; PGA, phosphoglycerate.

(Behrenfeld, M. J., Halsey, K. H. & Milligan, A. J. Phil Trans Royal Soc B: Biol Sci 363, 2687, 2008).

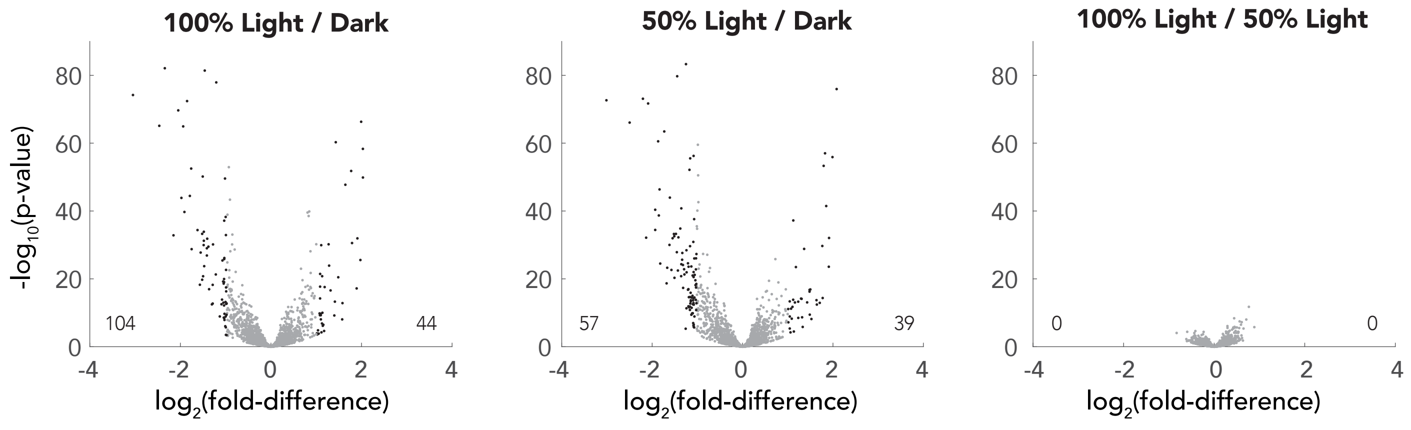


Figure S6. Effects of light exposure on bacterial gene expression as percent of transcriptome. Three light levels corresponding to those at the sample times in the co-culture experiment were examined: 100%, light level at noon; 50%, light level at mid-morning and mid-afternoon; dark, light level at night. Differentially expressed genes with fold-difference of $\geq$2 and DESeq2 adjusted-p $\leq$ 0.05 are shown as black symbols.

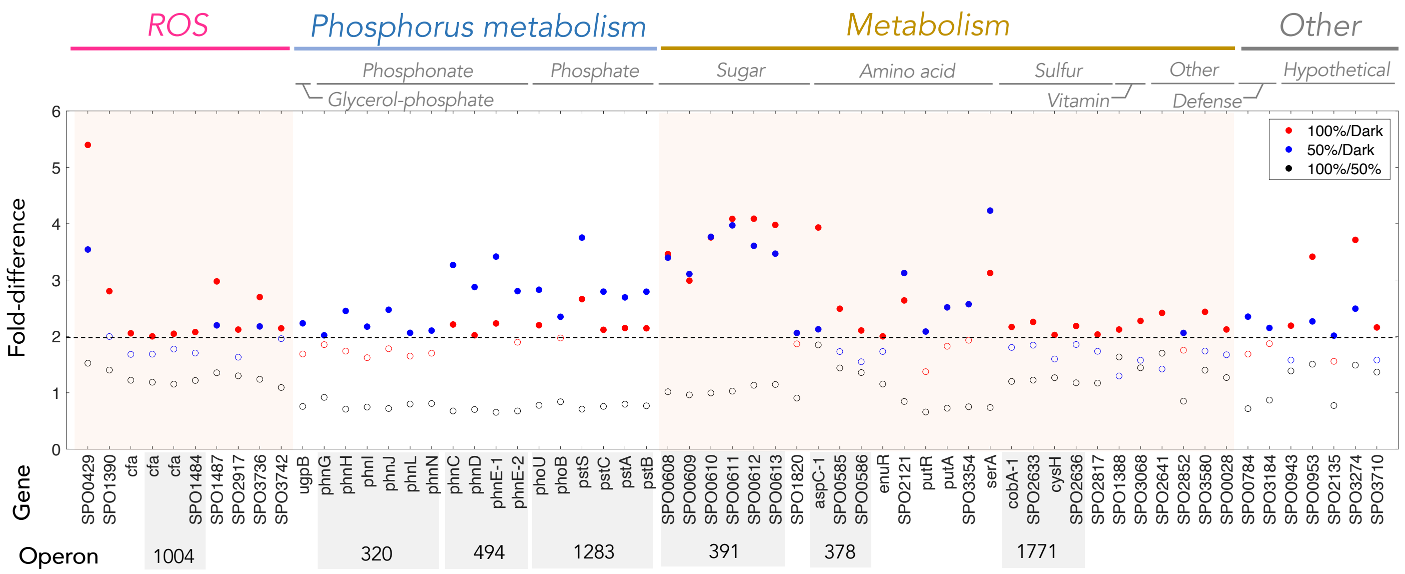


Figure S7. Direct effects of light on gene expression by *R. pomeroyi*. Filled circles indicate differentially expressed genes (fold-difference $\geq$2 and DESeq2 adjusted-p $\leq$ 0.05) at 100% light level relative to dark (red symbols), 50% light level relative to dark (blue symbols), and 100% light level relative to 50% light level (black symbols). Open circles with the same color codes represent comparisons that were not significantly different. All 61 genes enriched in the presence of light are included, and detailed gene information is given in Table S2. The horizontal line represents 2 fold-difference.

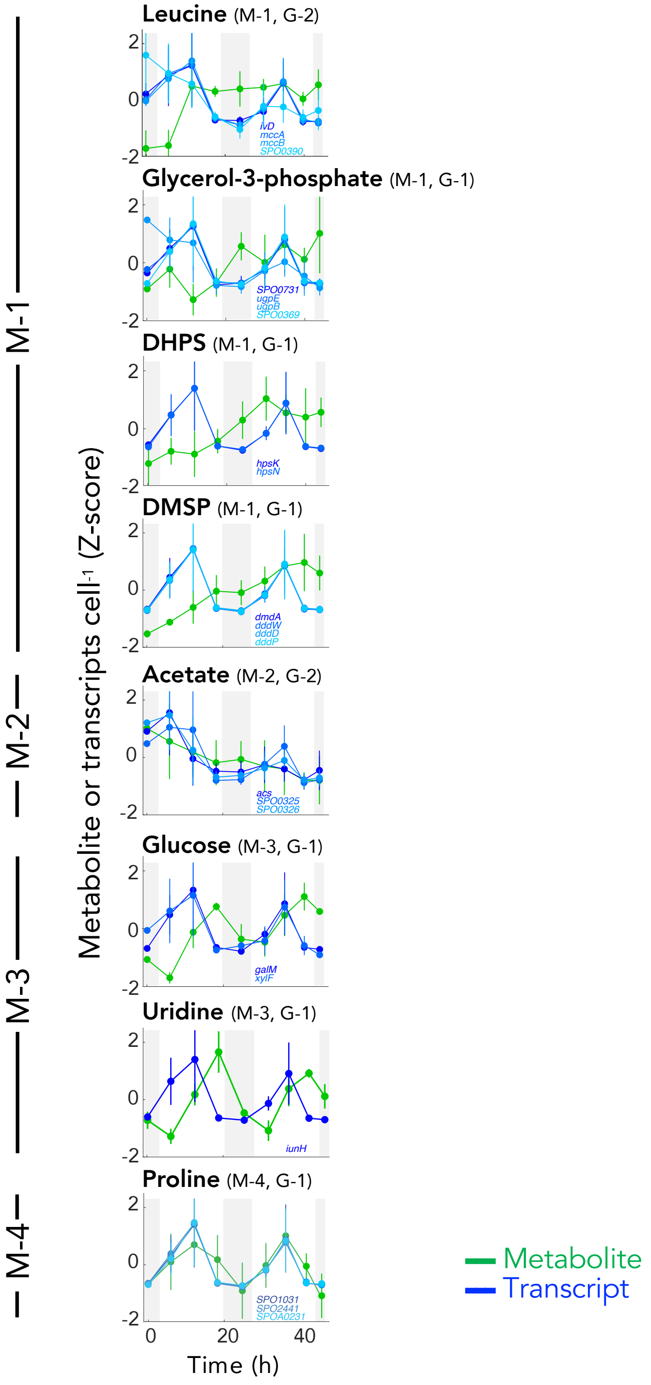


Figure S8. Temporal concentration changes in the eight diatom endometabolites (green) plotted with transcript inventories for all identified genes encoding uptake or catabolism of the same compound (blue shades). Error bars indicate standard deviation (n=3, except for genes in the first night where n=2).

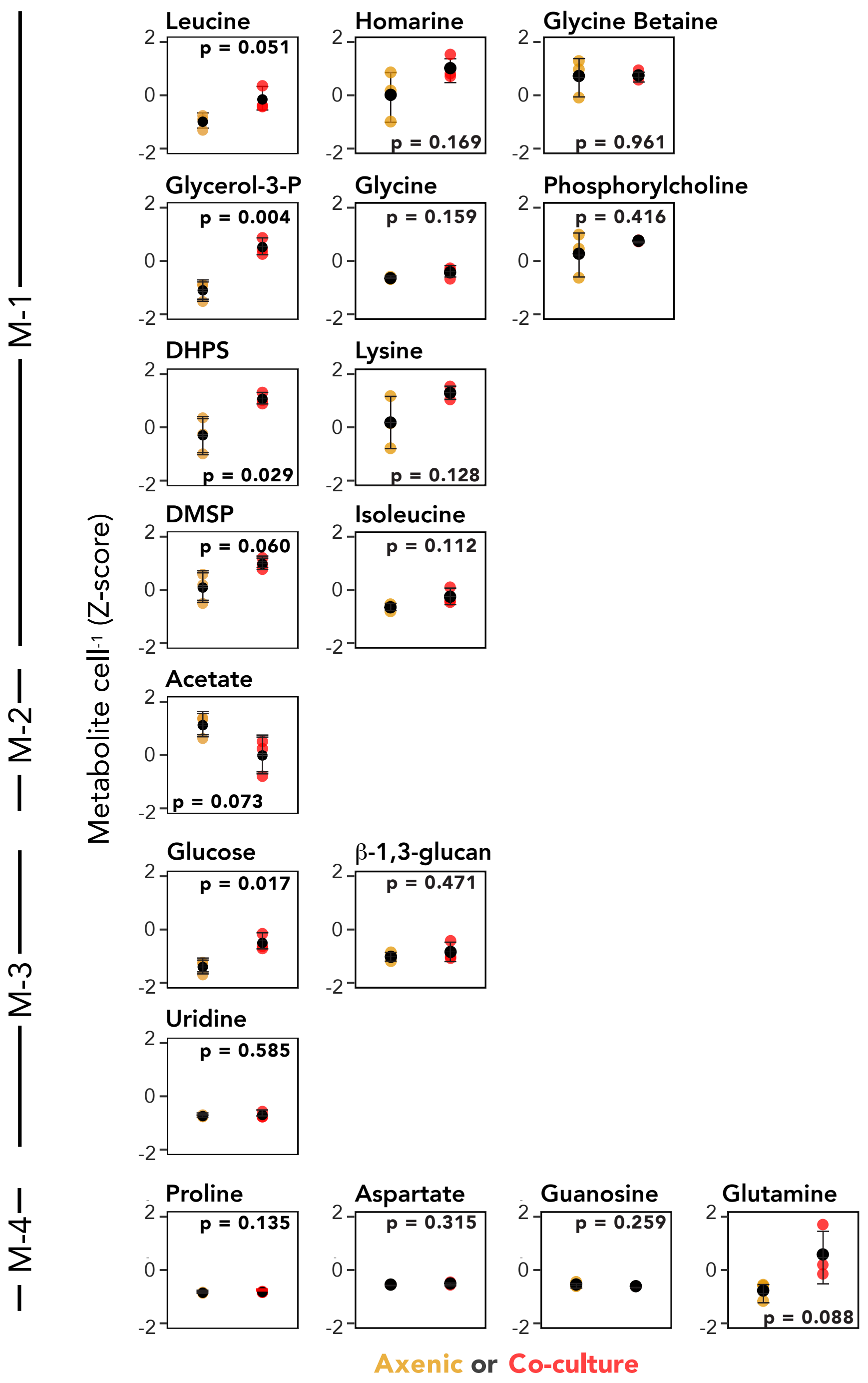


Figure S9. Comparison of endometabolite concentrations in axenic and co-culture treatments in the 15 day study. p values on each plot indicate results of t-tests. Black symbols indicate the mean value (n=3) and error bars indicate standard deviation.

Table S3. Summary of model output for the default and bacteria response models. Pearson correlations and *p* values between experimental and model data for metabolites and corresponding genes are shown. n.a., not applicable.

| Metabolite or gene name/locus tag | | Pattern category | Default model | | Recognition model | |
|---|---|---|---|---|---|---|
| | | | *r* | *p* | *r* | *p* |
| ***Metabolites*** | | | | | | |
| | Leucine | M-1 | -0.412 | 0.033 | 0.701 | 0.000 |
| | Glycerol 3-phosphate | M-1 | -0.181 | 0.365 | 0.554 | 0.003 |
| | DHPS | M-1 | -0.104 | 0.605 | 0.725 | 0.000 |
| | DMSP | M-1 | -0.246 | 0.217 | 0.824 | 0.000 |
| | Acetate | M-2 | 0.580 | 0.002 | n.a. | n.a. |
| | Glucose | M-3 | 0.807 | 0.000 | 0.913 | 0.000 |
| | Uridine | M-3 | 0.887 | 0.000 | n.a. | n.a. |
| | Proline | M-4 | 0.567 | 0.002 | n.a. | n.a. |
| ***Genes*** | | | | | | |
| *Leucine* | ivD | G-2 | 0.22 | < 0.05 | 0.17 | < 0.05 |
| | mccA | G-2 | 0.30 | < 0.01 | 0.21 | < 0.05 |
| | SPO0390 | G-2 | 0.01 | 0.73 | 0.07 | 0.18 |
| *Glycerol 3-phosphate* | SPO0731 | G-1 | 0.34 | < 0.01 | 0.26 | < 0.05 |
| | ugpE | G-1 | 0.32 | < 0.01 | 0.25 | < 0.05 |
| | SPO0369 | G-1 | 0.44 | < 0.001 | 0.35 | < 0.01 |
| *DHPS* | hpsK | G-1 | 0.42 | < 0.001 | 0.33 | < 0.01 |
| | hpsN | G-1 | 0.42 | < 0.001 | 0.34 | < 0.01 |
| *DMSP* | dmdA | G-1 | 0.42 | < 0.001 | 0.34 | < 0.01 |
| | dddW | G-1 | 0.45 | < 0.001 | 0.36 | < 0.01 |
| | dddD | G-1 | 0.42 | < 0.001 | 0.34 | < 0.01 |
| | dddP | G-1 | 0.44 | < 0.001 | 0.36 | < 0.01 |
| *Acetate* | SPO0325 | G-2 | 0.26 | < 0.05 | n.a. | n.a. |
| | SPO0326 | G-2 | 0.29 | < 0.01 | n.a. | n.a. |
| *Glucose* | galM | G-1 | 0.32 | < 0.01 | 0.31 | < 0.01 |
| | xylF | G-1 | 0.19 | < 0.05 | 0.17 | < 0.05 |
| *Uridine* | iunH | G-1 | 0.28 | < 0.05 | n.a. | n.a. |
| *Proline* | SPO1031 | G-1 | 0.25 | < 0.05 | n.a. | n.a. |
| | SPO2441 | G-1 | 0.24 | < 0.05 | n.a. | n.a. |
| | SPOA0231 | G-1 | 0.27 | < 0.05 | n.a. | n.a. |

Table S4. Instrument settings for NMR experiments.

| Experiment (program name by Bruker) | Corresponding figure and table | Spectral width/offset (ppm) | | Size of FID | | Number of scans |
|---|---|---|---|---|---|---|
| | | f2 | f1 | f2 | f1 | |
| $^{1}H$-$^{13}C$ HSQC (hsqcetgpprsisp2.2) | Figure 1, 3, S2, S8, and S9 | 14.2/4.7 | 40.0/68.0 | 1024 | 96 | 16 |
| | Fig. S1; Table S1 | 14.2/4.7 | 159.9/75.0 | 1024 | 256 | 256 |
| $^{1}H$-$^{13}C$ HSQC-TOCSY (hsqcdietgpsisp.2) | Fig. S1 | 14.2/4.7 | 189.9/90.0 | 1024 | 256 | 256 |
| $^{1}H$-$^{13}C$ HMBC (hmbcetgpl2nd) | Fig. S1 | 12.0/4.7 | 140/70.0 | 5768 | 390 | 96 |

Table S5. Summary of transcriptome sequence data. #reads, raw reads. #QC, number of reads passing quality control. QC%, percent of reads passing quality control. rRNA, number of reads identified as rRNA, %rRNA, percent of reads identified as rRNA, MTST5, number of reads identified as internal standard 5. MTST6, number of reads identified as internal standard 6. MTST_total, sum of MTST5 and MTST6 reads. std%, percent of reads identified as internal standards. #ReadsLeft, mRNA reads remaining for analysis. n.a., not applicable.

| ID | Sample | #reads | #QC | QC% | rRNA | rRNA% | MTST5 | MTST6 | MTST_total | std% | #ReadsLeft |
|---|---|---|---|---|---|---|---|---|---|---|---|
| E05 | Co-culture experiment, Night1-1 | 13,498,405 | 12,363,245 | 91.59 | 3,406,623 | 27.6 | 166,618 | 608,592 | 775,210 | 6.3 | 8,181,412 |
| F04 | Co-culture experiment, Night1-2 | 15,502,853 | 14,168,744 | 91.39 | 2,786,395 | 19.7 | 140,544 | 482,014 | 622,558 | 4.4 | 10,759,791 |
| F02 | Co-culture experiment, Mid-morning1-1 | 16,837,562 | 15,353,979 | 91.19 | 1,712,502 | 11.2 | 129,664 | 403,709 | 533,373 | 3.5 | 13,108,104 |
| E01 | Co-culture experiment, Mid-morning1-2 | 17,807,607 | 16,244,554 | 91.22 | 3,132,687 | 19.3 | 176,739 | 576,341 | 753,080 | 4.6 | 12,358,787 |
| E10 | Co-culture experiment, Mid-morning1-3 | 19,910,294 | 18,205,466 | 91.44 | 2,953,724 | 16.2 | 115,098 | 270,503 | 385,601 | 2.1 | 14,866,141 |
| F03 | Co-culture experiment, Noon1-1 | 19,275,856 | 17,722,932 | 91.94 | 1,164,010 | 6.6 | 115,767 | 324,058 | 439,825 | 2.5 | 16,119,097 |
| D08 | Co-culture experiment, Noon1-2 | 19,685,509 | 18,037,235 | 91.63 | 1,825,730 | 10.1 | 312,628 | 990,970 | 1,303,598 | 7.2 | 14,907,907 |
| E02 | Co-culture experiment, Noon1-3 | 19,408,658 | 17,720,533 | 91.30 | 966,890 | 5.5 | 81,101 | 241,754 | 322,855 | 1.8 | 16,430,788 |
| E11 | Co-culture experiment, Mid-afternoon1-1 | 20,885,480 | 19,054,538 | 91.23 | 3,581,342 | 18.8 | 324,456 | 959,900 | 1,284,356 | 6.7 | 14,188,840 |
| E12 | Co-culture experiment, Mid-afternoon1-2 | 17,557,260 | 16,073,027 | 91.55 | 3,914,892 | 24.4 | 261,021 | 719,276 | 980,297 | 6.1 | 11,177,838 |
| E03 | Co-culture experiment, Mid-afternoon1-3 | 16,977,882 | 15,523,460 | 91.43 | 1,406,118 | 9.1 | 348,476 | 1,057,642 | 1,406,118 | 9.1 | 12,711,224 |
| D12 | Co-culture experiment, Night2-1 | 23,851,789 | 22,079,516 | 92.57 | 7,073,693 | 32.0 | 233,505 | 1,014,268 | 1,247,773 | 5.7 | 13,758,050 |
| F07 | Co-culture experiment, Night2-2 | 19,469,653 | 17,836,736 | 91.61 | 3,663,942 | 20.5 | 197,821 | 734,821 | 932,642 | 5.2 | 13,240,152 |
| F09 | Co-culture experiment, Night2-3 | 31,934,931 | 29,269,343 | 91.65 | 5,338,425 | 18.2 | 479,384 | 1,609,432 | 2,088,816 | 7.1 | 21,842,102 |

| | | | | | | | | | | | |
|---|---|---|---|---|---|---|---|---|---|---|---|
| D10 | Co-culture experiment, Mid-morning2-1 | 20,554,827 | 18,824,484 | 91.58 | 2,744,962 | 14.6 | 186,059 | 571,690 | 757,749 | 4.0 | 15,321,773 |
| D11 | Co-culture experiment, Mid-morning2-2 | 13,279,747 | 12,174,740 | 91.68 | 1,927,329 | 15.8 | 203,698 | 551,681 | 755,379 | 6.2 | 9,492,032 |
| E04 | Co-culture experiment, Mid-morning2-3 | 14,605,987 | 13,367,467 | 91.52 | 2,215,170 | 16.6 | 109,203 | 347,448 | 456,651 | 3.4 | 10,695,646 |
| E06 | Co-culture experiment, Noon2-1 | 19,722,654 | 18,019,139 | 91.36 | 1,623,225 | 9.0 | 136,858 | 395,528 | 532,386 | 3.0 | 15,863,528 |
| E08 | Co-culture experiment, Noon2-2 | 28,885,375 | 26,482,283 | 91.68 | 1,079,553 | 4.1 | 117,326 | 323,826 | 441,152 | 1.7 | 24,961,578 |
| F06 | Co-culture experiment, Noon2-3 | 21,993,618 | 20,024,426 | 91.05 | 2,301,998 | 11.5 | 165,550 | 518,511 | 684,061 | 3.4 | 17,038,367 |
| E09 | Co-culture experiment, Mid-afternoon2-1 | 22,556,906 | 20,538,846 | 91.05 | 3,499,739 | 17.0 | 334,172 | 1,166,834 | 1,501,006 | 7.3 | 15,538,101 |
| F08 | Co-culture experiment, Mid-afternoon2-2 | 19,472,311 | 17,798,623 | 91.40 | 2,893,773 | 16.3 | 191,917 | 749,108 | 941,025 | 5.3 | 13,963,825 |
| D09 | Co-culture experiment, Mid-afternoon2-3 | 13,658,878 | 12,514,100 | 91.62 | 3,167,132 | 25.3 | 266,044 | 762,316 | 1,028,360 | 8.2 | 8,318,608 |
| E07 | Co-culture experiment, Night3-1 | 18,841,406 | 17,419,067 | 92.45 | 6,766,726 | 38.8 | 147,411 | 492,285 | 639,696 | 3.7 | 10,012,645 |
| F01 | Co-culture experiment, Night3-2 | 15,647,403 | 14,312,249 | 91.47 | 2,959,371 | 20.7 | 145,088 | 638,394 | 783,482 | 5.5 | 10,569,396 |
| F05 | Co-culture experiment, Night3-3 | 16,962,656 | 15,604,901 | 92.00 | 4,147,196 | 26.6 | 220,930 | 803,400 | 1,024,330 | 6.6 | 10,433,375 |
| 0-1 | Direct light experiment, light level 0%-1 | 25,586,505 | 24,300,490 | 94.97 | 6,613,045 | 27.2 | n.a. | n.a. | n.a. | n.a. | 17,687,445 |
| 0-2 | Direct light experiment, light level 0%-2 | 31,663,480 | 30,051,703 | 94.91 | 10,646,884 | 35.4 | n.a. | n.a. | n.a. | n.a. | 19,404,819 |
| 0-3 | Direct light experiment, light level 0%-3 | 26,282,421 | 24,902,417 | 94.75 | 9,821,337 | 39.4 | n.a. | n.a. | n.a. | n.a. | 15,081,080 |
| 50-1 | Direct light experiment, light level 50%-1 | 32,174,743 | 30,533,445 | 94.90 | 6,613,045 | 21.7 | n.a. | n.a. | n.a. | n.a. | 21,524,552 |
| 50-2 | Direct light experiment, light level 50%-2 | 29,926,762 | 28,405,710 | 94.92 | 10,646,884 | 37.5 | n.a. | n.a. | n.a. | n.a. | 17,997,524 |
| 50-3 | Direct light experiment, light level 50%-3 | 23,998,399 | 22,748,705 | 94.79 | 9,821,337 | 43.2 | n.a. | n.a. | n.a. | n.a. | 11,800,777 |

| | | | | | | | | | | | |
|---|---|---|---|---|---|---|---|---|---|---|---|
| 100-1 | Direct light experiment, light level 100%-1 | 26,526,300 | 25,152,623 | 94.82 | 6,613,045 | 26.3 | n.a. | n.a. | n.a. | n.a. | 15,731,031 |
| 100-2 | Direct light experiment, light level 100%-2 | 30,378,955 | 28,845,621 | 94.95 | 10,646,884 | 36.9 | n.a. | n.a. | n.a. | n.a. | 16,872,089 |
| 100-3 | Direct light experiment, light level 100%-3 | 33,568,541 | 30,855,170 | 91.92 | 9,821,337 | 31.8 | n.a. | n.a. | n.a. | n.a. | 19,179,745 |

**Modeling Methods**

To determine the factors impacting diel metabolite dynamics a simple model of the co-culture system was developed. The model consists of three metabolite pools, the phytoplankton endometabolome (*P*), the medium exometabolome (*E*), and the bacterial endometabolome (*B*). The time evolution of these pools was calculated using the following differential equations.

$$\delta_t P = N\,X - T - R$$

$$\delta_t E = R - U$$

$$\delta_t B = U - C$$

Where *N* is the metabolite biosynthesis rate, *X* is the bacterial response mechanism, increasing the metabolite biosynthesis rate in the presence of bacteria by 1.5- or 2-fold, *R* is release rate from the phytoplankton, and *T* is rate at which endometabolites are allocated for biomass and energy generation by phytoplankton cells. *U* represents bacterial uptake of the exometabolome and *C* represents catabolism of the metabolite within the bacterial metabolome. These differential equations were solved at times steps of 0.1 hours for 10 days using the variable definitions provided below. To simulate the experimental conditions the bacterial term was set to zero until inoculation on day 6.

*N* is represented by the following equation to account for the effects C-fixation irradiance oscillation around the peak in light intensity.

$$N = \begin{cases} -A\sin\left((t+0.5)*\frac{\pi}{12}+\frac{\pi}{6}\right)+\frac{1}{3}, & 0 < t \leq 16 \\ -0.5\,\mathrm{atan}(a_1 * t - a_1 * b_1) + 0.6, & 16 < t \leq (16+H) \\ -0.5\,\mathrm{atan}(a_2 * t - a_2 * b_2) + 0.6, & (16+H) < t \leq 24 \end{cases}$$

Where *A* represents the amplitude of *N*, *t* represents the independent variable time in hours, and *H* influences the intensity of this oscillation as the length of time, in tenths of an hour, it takes for *N* to fall to 30% efficiency. To align with the experiment's light-dark cycle the modeled light dark cycle is: dark from 0:00 hours to 08:00 hours, and light from 08:00 hours to 24:00 hours, with peak light intensity at 16:00 hours. The coefficients a and b were solved for as

$$a = \frac{\tan(-2N+1.2) - \tan(-2N_0+1.2)}{t - t_0}$$

$$b = \frac{-\tan(-2N+1.2)}{a} + t_0$$

so that the function *N* is continuous, where $N_0$ is the initial value of *N* for each time range and $t_0$ is the initial time. *T* was calculated as

$$T = P_0\,F_T$$

Where $P_0$ is the endometabolite concentration at the previous time point and $F_T$ is the fraction of the endometabolite consumed by internal cellular processes during each time step. $R$ was assumed to be release by passive diffusion out of the cell

$$R = F_R(P_0 - E_0 G)$$

Where $F_R$ is the fraction of the endometabolite pool that can be released from the cell in each time step and is multiplied by a term representing the diffusion gradient between endometabolome and the exometabolome at the previous time step, $E_0$. $G$ accounts for the difference in relative volume between the endometabolome and the exometabolome, relating the pool magnitude to a relative concentration. As an option within the base model $R_A$ allows the cell to increase release when light intensity is high, representing excretion for physiological balance:

$$R_A = a \cos\left(\frac{t\,\pi}{12} + \frac{8\pi}{12}\right) + 1$$

Where $a$ is the amplitude. $R_A$ then alters release rate as

$$R = F_R(P_0 - E_0 G)\, R_A$$

$U$ is assumed to follow Michaelis-Menten kinetics as

$$U = \frac{V_{max}\ E_0}{k_m\ +\ E_0}$$

Were $V_{max}$ is the maximum uptake velocity and $k_m$ is the half saturation constant. Finally, $C$ is represented as

$$C = B_0\, F_C$$

Where $B_0$ is the magnitude of the bacterial endometabolome at the previous time step and $F_C$ is the fraction of the bacterial endometabolite that is consumed in cellular processes each time step.

Table 1. Variable Model Parameters.

| | $P_0$ | $A$ | $F_T$ | $F_R$ | $G$ | $Vmax$ | $km$ | $H$ | $X$ | $R_A$ | $a$ |
|---|---|---|---|---|---|---|---|---|---|---|---|
| *M1G1Rec* | 20 | 0.05 | 0.002 | 0.001 | 0.05 | 0.67 | 3 | 30 | 2 | Yes | 0.5 |
| *M1G1* | 20 | 0.05 | 0.002 | 0.001 | 0.05 | 0.67 | 3 | 30 | 1 | Yes | 0.5 |
| *M1G2Rec* | 20 | 0.05 | 0.0015 | 0.0015 | 0.05 | 0.65 | 2 | 50 | 2 | Yes | 0.5 |
| *M1G2* | 20 | 0.05 | 0.0015 | 0.0015 | 0.05 | 0.65 | 2 | 50 | 1 | Yes | 0.5 |
| *M2G1* | 80 | 0.05 | 0.0015 | 0.0055 | 0.195 | 0.88 | 2 | 50 | 1 | Yes | 0.5 |
| *GlucRec* | 80 | 0.375 | 0.0015 | 0.001 | 0.05 | 0.8 | 2 | 70 | 2 | Yes | 0.5 |
| *M3G1* | 20 | 0.667 | 0.0055 | 0.003 | 0.05 | 0.74 | 2 | 79 | 1 | Yes | 1.0 |
| *M4G1* | 20 | 0.667 | 0.01 | 0.0085 | 0.05 | 0.75 | 2 | 20 | 1 | No | NA |
| *M4G2* | 20 | 0.6 | 0.0165 | 0.0165 | 0.05 | 0.39 | 1 | 21 | 1 | No | NA |
| *M2G2* | 80 | 0.25 | 0.0015 | 0.0035 | 0.15 | 0.7 | 6 | 65 | 1 | Yes | 1.0 |

$P_0$ is the initial phytoplankton endometabolite concentration.

$A$ is the amplitude of diel variation of metabolite synthesis.

$F_T$ is the fraction of the endometabolite pool consumed by internal cellular processes during each time step.

$F_R$ represents the baseline fraction of the endometabolite pool released from the cell in each time step.

$G$ influences the strengths of the diffusive gradient between the endometabolome and the exometabolome by accounting for the difference in relative volume, and subsequent concentration, between the pools.

$V_{max}$ is the maximum uptake velocity of exometabolite by bacteria.

$k_m$ is the half saturation constant of exometabolite uptake by bacteria.

$H$ influences the intensity of asymmetric carbon fixation as the length of time, in tenths of an hour, it takes for $N$ to fall to 30% efficiency.

$X$ is the optional bacterial recognition response, increasing the metabolite biosynthesis rate in the presence of bacteria.

$R_A$ represents increased excretion during high illumination for physiological balance, it is an option in the base model.

$a$ is the amplitude of the physiological release function, $R_A$.
